# FGF Signaling Regulates Development of the Anterior Fontanelle

**DOI:** 10.1101/2024.07.14.603452

**Authors:** Lauren Bobzin, Audrey Nickle, Sebastian Ko, Michaela Ince, Arshia Bhojwani, Amy E. Merrill

## Abstract

The calvarial bones of the infant skull are connected by transient fibrous joints known as sutures and fontanelles, which are essential for reshaping during birth and postnatal growth. Genetic disorders such as Apert, Pfeiffer, Crouzon, and Bent bone dysplasia linked to *FGFR2* variants often exhibit multi-suture craniosynostosis and a persistently open anterior fontanelle (AF). This study leverages mouse genetics and single-cell transcriptomics to determine how *Fgfr2* regulates closure of the AF closure and its transformation into the frontal suture during postnatal development. We find that cells of the AF, marked by the tendon/ligament factor SCX, are spatially restricted to ecto- or endocranial domains and undergo regionally selective differentiation into ligament, bone, and cartilage. Differentiation of SCX+ AF cells is dependent on FGFR2 signaling in cells of the osteogenic fronts which, when fueled by FGF18 from the ectocranial mesenchyme, express the secreted WNT inhibitor WIF1 to regulate WNT signaling in neighboring AF cells. Upon loss of *Fgfr2*, *Wif1* expression is lost, and cells of the AF retain a connective tissue-like fate failing to form the posterior frontal suture. This study provides new insights into regional differences in suture development by identifying an FGF-WNT signaling circuit within the AF that links frontal bone advancement with suture joint formation.

## Introduction

The calvarial bones of the infant skull are linked by transient fibrous joints called sutures and fontanelles. Sutures, which join two adjacent calvarial bones, and fontanelles, which occupy large, unossified regions between multiple bones where sutures will eventually form, are critical for calvarial reshaping during birth and postnatal calvarial growth. Numerous genetic disorders present with craniofacial deformities when sutures fuse prematurely (craniosynostosis), or fontanelles remain patent. For example, syndromes associated with genetic variants in Fibroblast growth factor receptor 2 (*FGFR2)* including Apert, Pfeiffer, Crouzon, and Bent bone dysplasia often present with multi-suture craniosynostosis, as well as a persistently open anterior fontanelle (Azoury et al., 2017, Merrill et al., 2012, Reardon et al., 1994, Wilkie et al., 1995). The co-occurrence of these seemingly opposite calvarial joint phenotypes suggests important differences in the way FGF signaling regulates development of sutures versus fontanelles.

The anterior fontanelle (AF), colloquially known as a baby’s “soft spot”, is a broad area of fibrous connective tissue that forms at the apex of the embryonic calvaria. It is here that the two frontal and two parietal bones will eventually meet at the bregma. The rapid apical expansion of these bones occurs during postnatal development via intramembranous ossification, replacing the AF with the posterior frontal suture (PFS) which joins the paired frontal bones from the bregma to the jugum limitans. Once the PFS is established, it stratifies into two distinct layers of bone, with the endocranial layer undergoing fusion through endochondral-like ossification which requires *Sox9* and regional inhibition of WNT signaling (Bradley et al., 1996, Behr et al., 2013, Manzanares et al., 1988, Sahar et al., 2005). The fate of the AF connective tissue and its role in frontal suture development, as well as its potential contribution to the etiology of calvarial defects, has remained unclear.

While calvarial phenotypes in *FGFR2*-related syndromes indicate a key role for FGF signaling in the developing AF, studies to date have largely focused on the receptor’s role in bone formation during embryogenesis when it is expressed within the mid-suture and advancing osteogenic bone fronts. Genetic studies in mice have shown that FGFR2 signaling, likely activated by FGF18, is critical to maintain the balance of proliferation and osteoblast differentiation within sutures (Ohbayashi et al., 2002). In mouse models that harbor gain-of-function *Fgfr2* variants associated with human craniosynostosis syndromes, mechanisms of premature fusion of the coronal suture include altered proliferation and induction of ectopic osteoblast differentiation (Holmes et al., 2009, Wang et al., 2005, Yin et al., 2008, Pfaff et al., 2016, Hajihosseini et al., 2001). These mouse models also exhibit a midline gap between the frontal bones in a region anatomically homologous to the human AF. Studies using the Apert syndrome mouse model have shown that this midline gap is linked to decreased osteoblast differentiation, as well as transcriptomic changes in the cells within the osteogenic fronts of the frontal bones (Wang et al., 2005, Holmes et al., 2020). While a midline ossification defect was also reported in mice with conditional loss of *Fgfr2* in skeletogenic mesenchyme, the mechanism underlying the phenotype has yet to be investigated (Yu et al., 2003).

Here, we show that *Fgfr2* is necessary for AF closure and, subsequently, frontal suture joint formation. We find that cells of the AF express the ligament marker *Scx* and represent at least two transcriptionally distinct populations spatially restricted to either the ecto- or endocranial domains, which give rise to bone and cartilage in an FGF-dependent manner. We provide evidence that FGF signaling in the osteogenic fronts promotes expression of the secreted WNT inhibitor WIF1 that non-autonomously regulates WNT signaling in AF cells. In the absence of FGF signaling, *Wif1* expression is lost, and cells of the AF fail to differentiate into bone and cartilage, instead maintaining a connective-tissue like identity. Correspondingly, elevation of WNT/β-catenin signaling induces a persistent AF phenotype and downregulation of WNT/β-catenin rescues AF closure in *Fgfr2* mutant mice. Together, these results suggest that FGF-mediated regulation of WNT signaling regionally influences cell fate and suture formation within the AF.

## Results

### *Fgfr2* is necessary for closure of the anterior fontanelle

Genetic lineage analysis in mice using *Wnt1-Cre* previously showed that neural crest-derived mesenchyme gives rise to the frontal bones, as well as the tissue that occupies the AF (**Figure 1A**) (Jiang et al., 2002, Yoshida et al., 2008). Therefore, to investigate a role of FGFR2 within the AF development, we performed *Wnt1-Cre*-mediated deletion of *Fgfr2* using a floxed allele that produces a functional null copy of the receptor (Yu et al., 2003). Since ablation of *Fgfr2* occurs in neural crest cells (NCC) and their derivatives, we will refer to them as NCC-*Fgfr2^-/-^* mice. We examined calvarial bone and cartilage development in NCC-*Fgfr2^-/-^*and littermate controls using wholemount alizarin red and alcian blue staining. The AF of NCC-*Fgfr2^-/-^* mice is wide and patent beginning at embryonic day (E) 18.5 compared to control (**Figure 1B,E**). In control mice at postnatal day (P) 3, the AF begins to progressively close, while that of NCC-*Fgfr2 ^-/-^* mice remains wide open (**Figure 1C,F**). At P5, the frontal bones of control mice have closely approximated to form the PFS, while the AF remains persistent in the mutant (**Figure 1D,G**). Quantitative measurement showed that the normalized average area between the frontal bones (total area of AF/length of PFS from jugum limitans to bregma) is significantly larger in NCC-*Fgfr2^-/-^* at P3 (p=.0.0167, n=7) and P5 (p=.0003, n=9) compared to littermate controls (**Figure 1H**). By P10, the PFS of control mice begins to form cartilaginous fusions that follow a joint-gap pattern, while the AF has failed to resolve into the PFS in NCC-*Fgfr2^-/-^*mice (**Figure 1I,L**). MicroCT analysis shows that the PFS of control mice has fused by P30, with distinct, bony endocranial and ectocranial layers visible in orthoslice views (**Figure 1J,K**). In NCC-*Fgfr2^-/-^*mice, the AF persists while regions of the frontal and sagittal sutures that it prefigures fail to form (**Figure 1M,N**). Together these results show that *Fgfr2* is necessary for AF closure.

**Figure 1.**
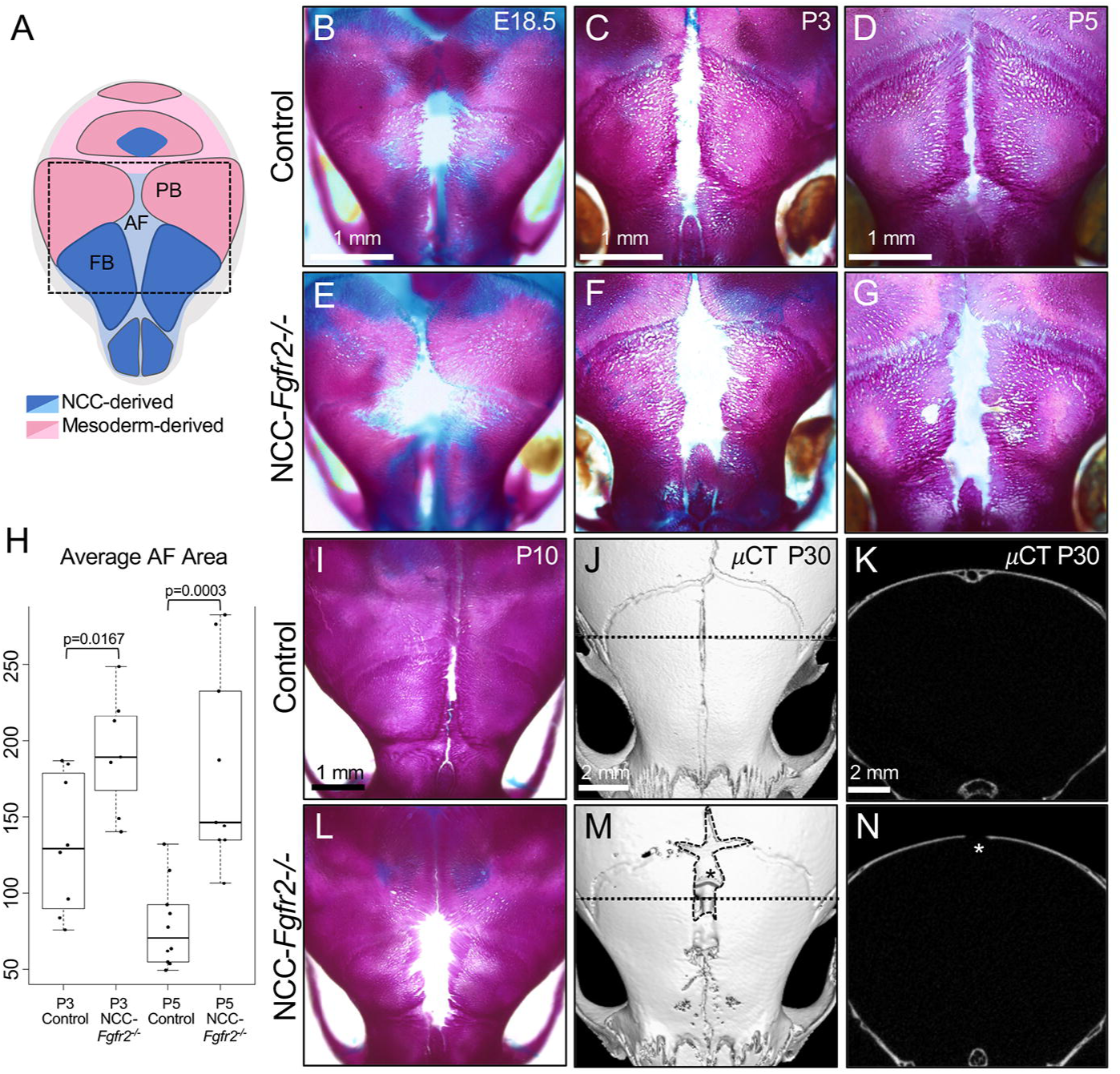
*Fgfr2* is required in NCC-derived mesenchyme for closure of the AF and formation of PFS. (A) Diagram representing the tissue origins of the calvarial vault. NCC-derived regions are depicted in blue and paraxial mesoderm-derived regions are depicted in pink. Darker shades of either color represent bone while lighter shades represent fibrous connective tissue, including the fontanelle. Wholemount skeletal prep staining using alizarin red (bone) and alcian blue (cartilage) shows morphological differences between the AF of controls and NCC-*Fgfr2^-/-^*littermates throughout development. (B,E) At E18.5, NCC-*Fgfr2^-/-^*mice show a slightly wider anterior fontanelle compared to control. (C-D,F-G) By P3 and P5, the difference becomes more pronounced as the frontal bones of NCC-*Fgfr2^-/-^* mutants fail to approximate. (H) Quantification of the averaged normalized AF area at P3 and P5 shows statistically significant differences between control and NCC-*Fgfr2^-/-^*mice. (I,L) At P10, the control PFS has begun to fuse anteriorly, forming cartilage adjacent to the jugum limitans, while NCC-*Fgfr2^-/-^* mutants completely lack this cartilage as frontal bones still fail to advance. (J-K,M-N) µCT scans of NCC-*Fgfr2^-/-^* mutant and littermate control at P30 show persistency of the AF at the bregma (marked with asterisk) and dysmorphia of the anterior PFS. Dotted lines in J and M correspond to orthoslices that highlight the complete lack of frontal bone fusion and suture organization (asterisk; K,N). AF, anterior fontanelle; PFS, posterior frontal suture; PB, parietal bone; FB, frontal bone. N = at least 7 littermate pairs for each stage.

### Loss of *Fgfr2* blocks poster frontal suture formation within the anterior fontanelle

The posterior frontal suture, which forms within the anterior fontanelle, undergoes normal fusion in the endocranial domain through a cartilage intermediate. To identify histogenic changes in PFS development Hall-Brunt Quadruple stain (HBQ) was used to distinguish bone and cartilage (Hall, 1986). In control mice at P5, the paired frontal bones have approximated in the ectocranial domain and secondary osteogenic fronts have formed within the endocranial domain (**Figure 2A**). The frontal bones of NCC-*Fgfr2^-/-^* mice, however, remain far apart and the secondary osteogenic fronts within the endocranial domain are missing (**Figure 2B**). In control mice at P7, the cartilage template that prefigures the fused endocranial layer of the PFS begins to condense, becoming more differentiated at P11 and P13 (**Figure 2C,E,G**). Conversely, the frontal bones of NCC-*Fgfr2^-/-^*mice remain separated and show no evidence of cartilage formation or layered stratification (**Figure 2D,F,H**). All together these results show that *Fgfr2* is necessary for formation and fusion of the PFS within the AF.

**Figure 2.**
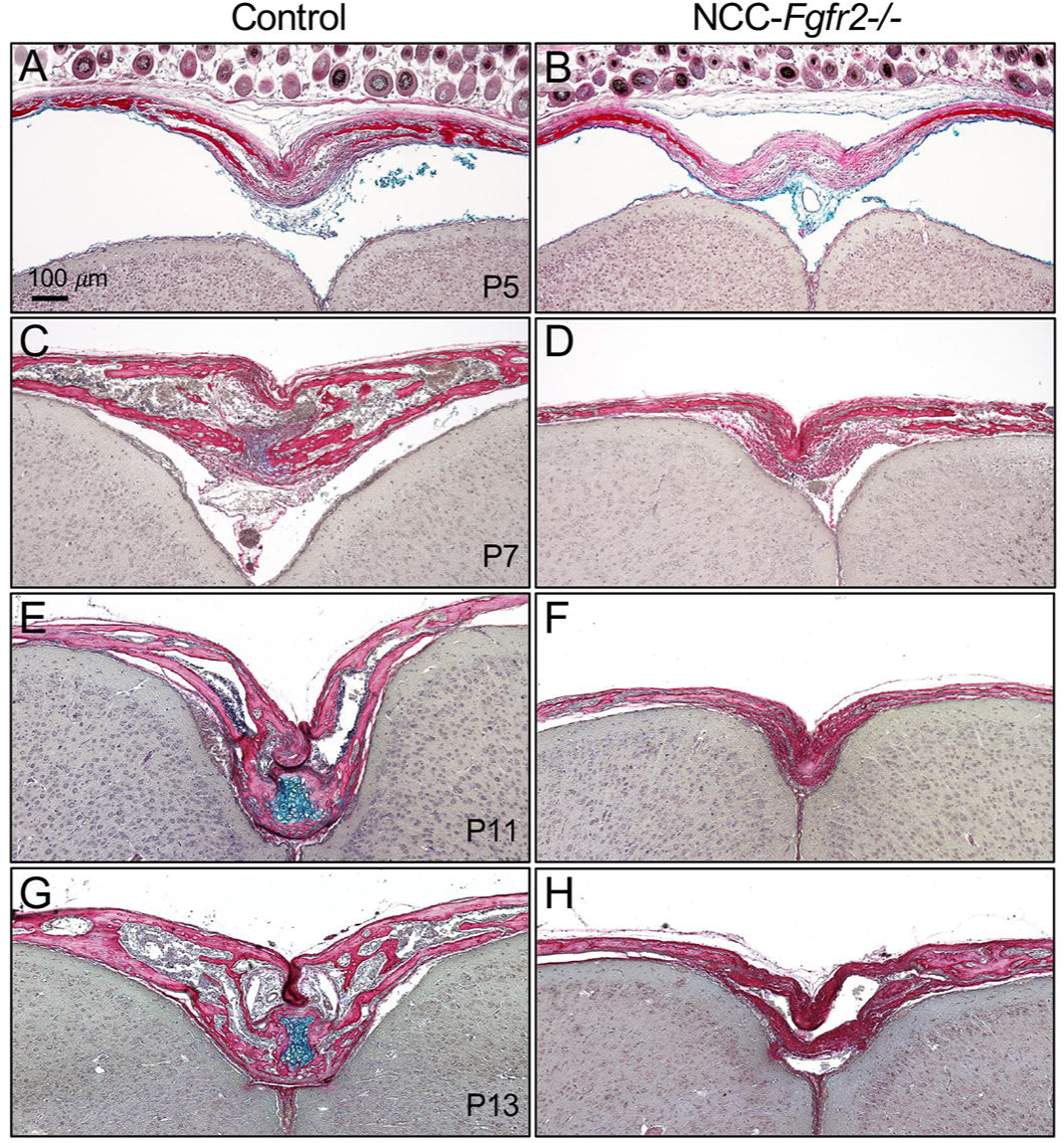
*Fgfr2* is required in NCC-derived mesenchyme for closure of the AF. HBQ stained coronal sections at critical stages of PFS development. (A,B) At P5 the ectocranial and endocranial layers AF are less defined and the frontal bones are further apart in NCC-*Fgfr2^-/-^* mutants compared to littermate controls. (C,D) At P7, when cartilage begins to condense in the endocranial layer in control, there is no evidence of cartilage in NCC-*Fgfr2^-/-^* mice. (E-H) At P11 and P13, when the endocranial cartilage matures and begins to ossify in control, there remains no cartilage in NCC-*Fgfr2^-/-^* mutants and the frontal bones remain separated in both the ecto- and endocranial domains by connective tissue. N = at least 4 littermate pairs per stage.

### *Fgfr2* regulates cell identity in the anterior fontanelle

To better understand *Fgfr2*+ cell populations in the developing AF, we analyzed a previously published single cell RNA sequencing (scRNA-seq) dataset of the developing frontal suture at E18.5, which includes the paired osteogenic fronts of the frontal bones and intervening AF connective tissue (FaceBase data set 1-4TT6) (Holmes et al., 2020). Re-analysis of this dataset using Seurat identified 9 mesenchymal clusters with similar gene enrichment to those previously reported (**Figure 3A,C**) (Holmes et al., 2020). Violin plots show *Fgfr2* expression is enriched in cell clusters we designated anterior fontanelle (AF) 1 and 2 based on their expression of genes associated with ligament-like connective tissue including *Scx*, *Tnmd*, *Mkx*, and *Matn4* (**Figure 3B**). Interestingly, AF2 is distinct from AF1 due to its enriched expression of the chondrogenic gene *Sox9* and tenogenic gene *Egr1* (**Figure 3B**). This is consistent with a previous study showing that *Sox9* is indispensable for closure of the PFS through endochondral-like ossification (Sahar et al., 2005). *Fgfr2* expression is most enriched in the osteogenic populations including the osteoblasts (OB) cluster marked by *Ibsp* expression and osteogenic front (OF) 1 and 2 clusters that express pro-osteogenic genes *Npnt* and *Sp7* (**Figure 3B**). Interstingly, *Sp7* has been implicated in regulating cellular recruitment at advancing bone fronts (Kague et al., 2016, Wang et al., 2022). Compared to OF1, the OF2 cluster is enriched for AF2-related genes including *Sox9* and *Egr1*. Relatively lower levels of *Fgfr2* are expressed in the dura mater (DM) cluster, which is marked by expression of *Cxcl12* (**Figure 3B**) (Holmes et al., 2020). While the *Fgfr2* expression is not enriched in the ectocranial mesenchyme (EM) cluster marked by *C1qtnf3* and *Igfbp3* or the hypodermis (HD) cluster marked by *Clec3b,* these clusters express the FGFR2 ligand *Fgf18* (**Figure 3B**) (Farmer et al., 2021). RNAScope fluorescent *in situ* hybridization at E18.5 validates that the spatial distribution of *Fgfr2* expression overlaps domains for the AF1 and AF2 marker *Tnmd* and the OF1 and OF2 marker *Sp7* (**Figure 3D,E**). Pseudotime analysis performed with Monocle 3 predicts that the AF1 and AF2 clusters followed distinct developmental trajectories to give rise to OF1 and OF2, respectively, and converge on an osteogenic identity (**Figure 3F**). This likely represents the distinct contribution of *Sox9*+ AF2 cells to endochondral-like bone that forms within the endocranial domain of the PFS. Cellular communication modeling using CellChat indicates that indicates that FGF18 produced by the EM is the critical source of ligand for FGFR2 expressing cells in the AF and OF populations (**Figure 3G,H**)(Jin et al., 2021).

**Figure 3.**
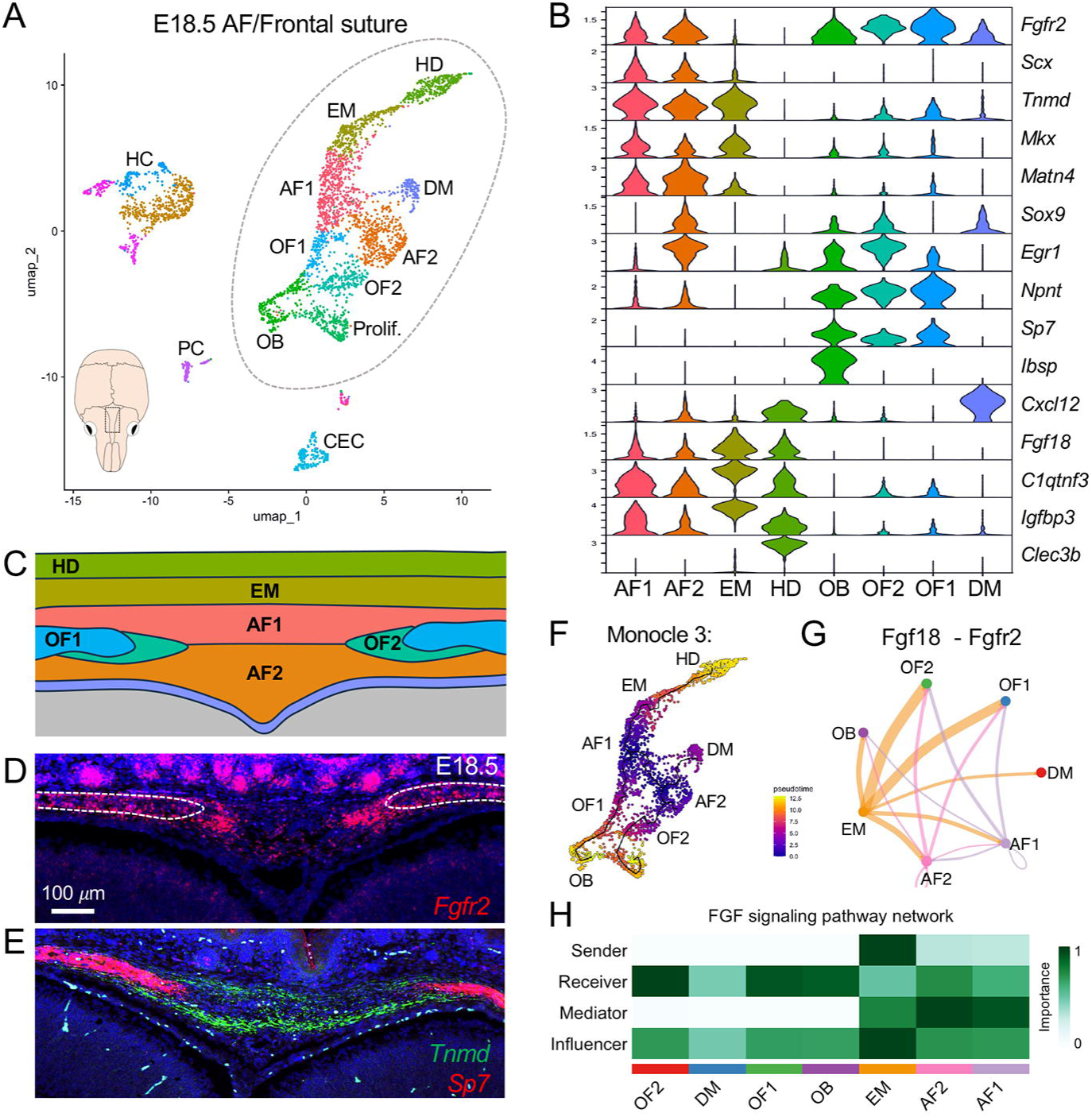
Single cell RNA-seq identifies heterogenous cell populations within the developing AF. (A) UMAP visualization for single cell RNA-sequencing of the E18.5 frontal suture. Mouse skull diagram includes boxed region highlighting the tissue collected for analysis. Dotted circle outlines mesenchymal cell types which are more closely delineated in our analysis. (B) Violin plots show mesenchymal clustered enriched for *Fgfr2* expression along with cluster specific gene expression. (C) Diagram of the developing AF showing precise locations of each mesenchymal cell cluster based on our analysis and a previous study that described this dataset (Holmes et al., 2020). Colors coordinate with cluster identities shown in panels A and B. (D,E) *In situ* hybridization validates expression of *Fgfr2* (red) in OF1 and OF2, that are marked by *Sp7* (red), and AF1 and AF2 that are marked by *Tnmd* (green) in the AF at E18.5. Dotted lines in panel D outline osteogenic fronts. (F) Pseudotime analysis performed with Monocle 3 predicts that cells in AF1 and AF2 clusters differentiate into OF1 and OF2, respectively. (G,H) Cell chat analysis predicts that the ectocranial mesenchyme is a source of FGF18 that signals to *Fgfr2*-expressing cells in the osteogenic front and anterior fontanelle, with the osteogenic cells (OF1, OF2, and OB), being the primary receivers. AF, anterior fontanelle; CEC, capillary endothelial cells; DM, dura mater; EM, ectocranial mesenchyme; HC, hematopoietic cells; HD, hypodermis; OB, osteoblasts; OF, osteogenic front; PC, pericytes; Prolif., proliferating osteoblasts.

To determine how loss of *Fgfr2* impacts the cell populations identified by scRNA-seq, we examined changes in cluster-enriched gene expression in the developing AF at E18.5 using RNAScope *in situ* hybridization. In the control, *Scx*, which marks the AF1 and AF2 clusters, is expressed in the anterior fontanelle connective tissue between the bone fronts with highest expression in the ectocranial region (**Figure 4A**). The expression domain of *Scx* expression is expanded laterally in NCC-*Fgfr2^-/-^*littermates, likely indicative of the increased size of the AF in mutants (**Figure 4B**). *Sp7*, a marker for the OF1 and OF2 clusters, is expressed in osteogenic fronts of controls and reduced in NCC-*Fgfr2^-/-^* littermates (**Figure 4C,D**). In control, *Sox9*, a marker for the AF2 and OF2 clusters, is expressed in the endocranial region of the AF and within the osteogenic fronts (**Figure 4C,C’**). Expression of *Sox9* is reduced in the AF and osteogenic fronts of NCC-*Fgfr2^-/-^* littermates (**Figure 4D,D’**). This suggests that loss of *Fgfr2* leads to a reduction of cells with the AF2 and OF2 identity. On the other hand, no difference in the expression of *Fgf18*, a marker for the EM cluster, was detected (**Figure 4E,F**). Together these results provide strong evidence that transcriptionally distinct populations of AF cells occupy spatially separate domains and are differentially impacted by loss of FGFR2 signaling.

**Figure 4.**
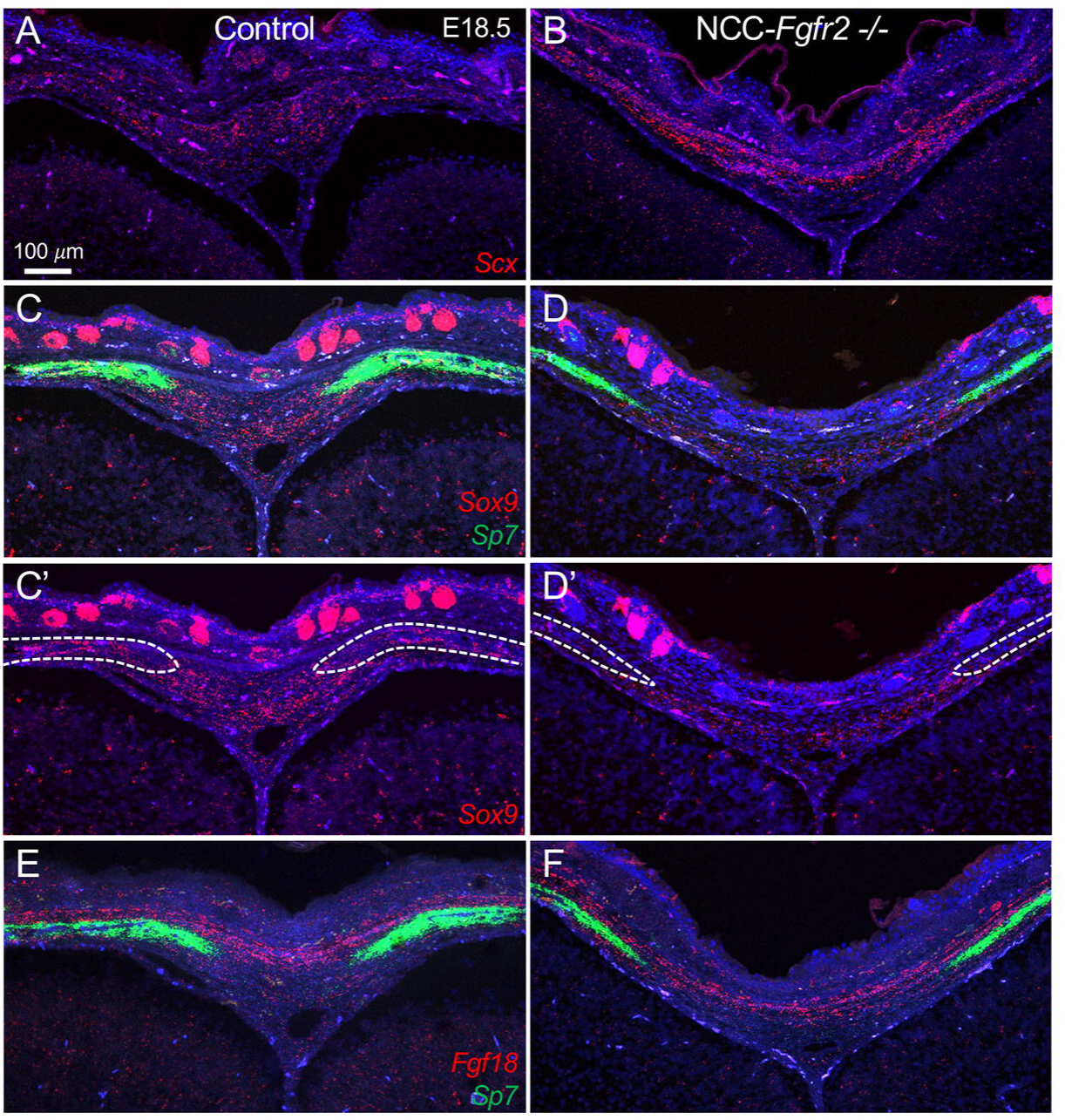
Expression of cell-type specific genes are altered in AF of NCC-*Fgfr2^-/-^* embryos. *In situ* hybridization with RNAScope was used to identify changes in cluster specific gene expression in the AF at E18.5. (A,B) *Scx*, a marker for AF1 and AF2 is expressed within the AF in control. This expression domain is expanded laterally towards the osteogenic fronts in the NCC-*Fgfr2^-/-^* mutant AF. (C,C’) In control, *Sox9*, a marker for AF2 and to a lesser extent OF2, is localized to the endocranial domain of the AF and within the bone. *Sp7*, a marker for OF1 and OF2, is expressed throughout the bone. (D,D’) In the NCC-*Fgfr2^-/-^* AF, *Sox9* expression is reduced within the AF and largely missing from the bone labelled with *Sp7*. (E,F) The domain of *Fgf18* expression in the EM shows little to no difference between control and NCC-*Fgfr2^-/-^*mutant AF relative to the osteogenic fronts marked by *Sp7* expression. Lines in panels C’ and D’ indicate the domain of *Sp7* expression in panels C and D. N = 3 littermate pairs per stage.

### *Fgfr2* is required for osteogenic differentiation of anterior fontanelle cells

Very little is known about the AF connective tissue cells and their role in frontal suture formation. To observe AF connective tissue cells during calvarial development, we employed a *Scx-GFP* transgenic reporter mouse that has been previously used to study tendon and ligament development (Pryce et al., 2007). At E18.5, Scx-GFP was expressed in all calvarial sutures and fontanelles, as well as immediately adjacent bone (**Figure 5A,C**). At P3, when the frontal suture has largely replaced the AF, Scx-GFP expression is confined to the suture connective tissue (**Figure 5B,D**). Sections through the AF at these stages reveals a bi-layered expressional pattern of Scx-GFP representing the ecto- and endocranial layers of the suture mesenchyme, as well as some GFP+ cells observed within the frontal bones at P3 (**Figure 5E,F**). There is notably increased Scx-GFP expression in the ectocranial layer which will eventually establish the permanent ligament of the suture.

**Figure 5.**
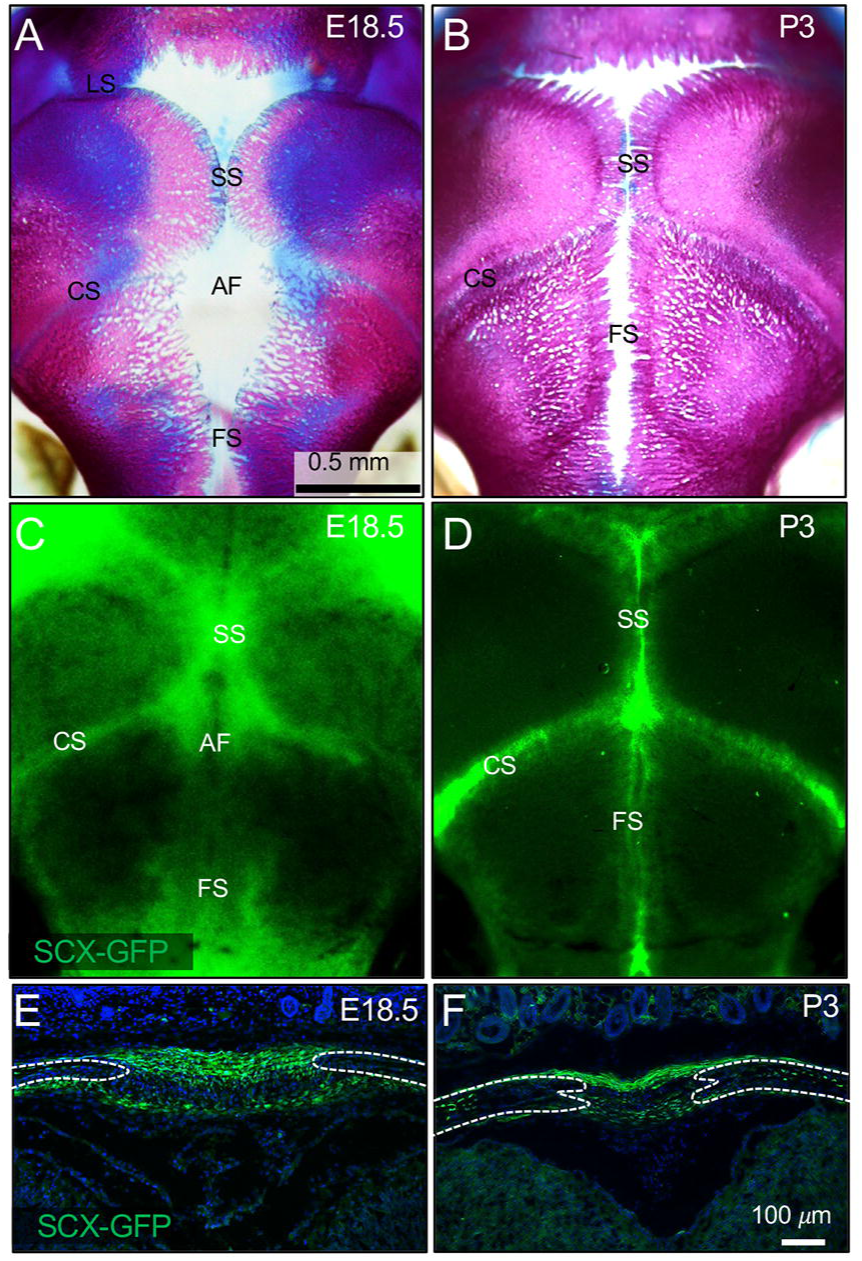
ScxGFP, a reporter for tendon/ligament, is expressed in the AF and PFS. (A,B) Wholemount alizarin red and alcian blue staining of the calvaria at E18.5 and P3 is shown to discern bone and cartilage. (C) At E18.5, Scx-GFP is expressed throughout the connective tissue of AF, as well as in the coronal and sagittal sutures. (D) At P3, when the AF is being replaced by the PFS, Scx-GFP expression is more confined to the sutures themselves. (E,F) Coronal sections through the anterior fontanelle at E18.5 and P3 show a bi-layered expressional pattern of Scx-GFP representing the ecto- and endocranial layers. Osteogenic fronts are outlined with dotted lines. AF, anterior fontanelle; CS, coronal suture; FS, frontal suture; SS, sagittal suture.

We next examined osteogenic differentiation within the AF of NCC-*Fgfr2^-/-^*mice using immunofluorescent detection of RUNX2, an early marker of osteoblasts, along with Scx-GFP to discriminate the AF connective tissue. At P0, RUNX2+ cells are enriched within the osteogenic fronts of the paired frontal bones, while Scx-GFP expression is localized to cells in the ecto- and endocranial layers of the AF in both control and NCC-*Fgfr2^-/-^* mice (**Figure 6A,B**). In control mice at P3, RUNX2 expression is activated in SCX+ cells located in the endocranial layer of the AF connective tissue, and SCX+/RUNX2+ double positive cells are found in the advancing frontal bones (**Figure 6C**). SCX+/RUNX2+ cells fail to form in the endocranial layer of NCC-*Fgfr2^-/-^* mice remaining exclusively SCX+ within the AF (**Figure 6D**). The ectocranial layer of SCX+ cells spanning and overlaying the frontal bones at this remained similar between control and mutant mice (**Figure 6C,D**). In controls at P5, RUNX2+ cells coalesce at the approximating endocranial osteogenic fronts to form the PFS, and SCX+/RUNX2+ cells show contribution to the approximating bone fronts. Meanwhile, the ectocranial layer in control mice is still occupied by SCX+ connective tissue **(Figure 6E)**. Conversely, in NCC-*Fgfr2^-/-^* mice the AF remains patent and very few SCX+/RUNX2+ are detected in the osteogenic fronts or bones (**Figure 6F**).

**Figure 6.**
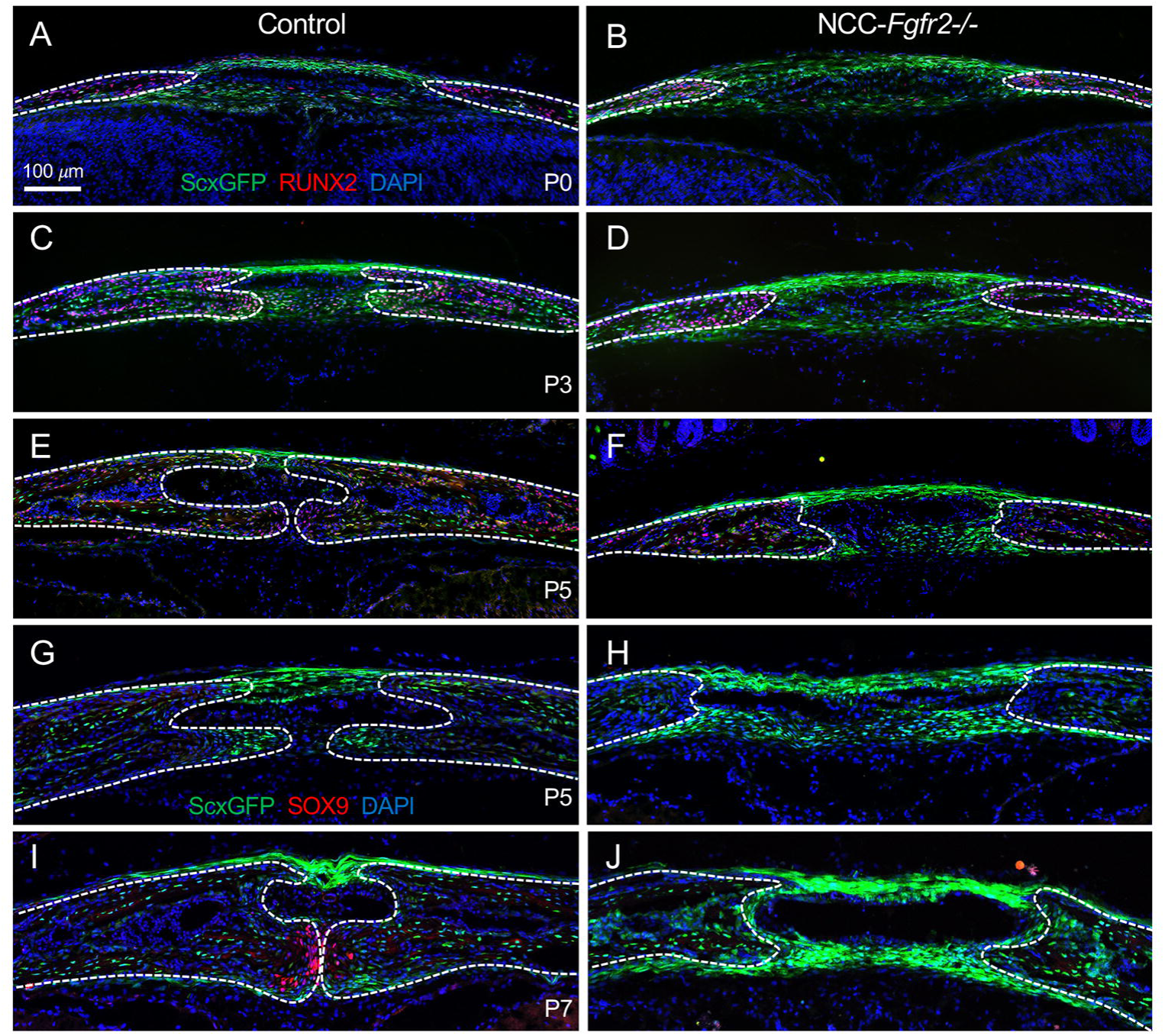
*Fgfr2* is necessary for differentiation of SCX+ cells of the AF into skeletogenic cells expressing RUNX2 and SOX9 during PFS formation. (A,B) Scx-GFP (green) expression along with immunofluorescence labeling of preosteoblast marker RUNX2 (red) shows that SCX+ cells within the AF are non-osteogenic in control and NCC-*Fgfr2^-/-^* mice at P0. (C,D) In control at P3, during PFS formation, SCX+ cells within the AF begin to express RUNX2 (SCX+/RUNX2+). RUNX2+ remains undetectable in SCX+ cells within the NCC-*Fgfr2^-/-^* AF. (E,F) In control at P5, after the PFS is established, most SCX+ cells in express RUNX2 with the exception of SCX+ cells just above the suture that will form the suprasutural ligament. (G,H) Scx-GFP (green) expression along with immunofluorescence labeling of the chondrocyte marker SOX9 (red) identified no SOX9+ cells in the PFS of control or NCC-*Fgfr2^-/-^*mice at P5. (I,J) At P7, while SCX+ cells begin expressing SOX9+ (SCX+/SOX+) within the endocranial domain of the control PFS, SOX9+ cells fail to form in the NCC-*Fgfr2^-/-^* PFS. Dotted lines in each panel outline osteogenic fronts. N = 3 littermate pairs per stage.

To detect chondrogenic differentiation, we performed immunofluorescent detection of SOX9, an early marker for chondrocytes. While SOX9 is not detected in the PFS at P5 in control or NCC-*Fgfr2^-/-^*mice (**Figure 6G,H**), a population of SOX9+ cells appear in the endocranial region of the suture midline in controls by P7 (**Figure 6I**). At this stage, the SCX+ suprasutural ligament is clearly distinguished (**Figure 6I**). In NCC-*Fgfr2^-/-^*mice, however, SOX9 expression remains undetectable, and the persistent AF is occupied by SCX+ cells that structurally resemble suprasutural ligament in both the ecto- and endocranial layers (**Figure 6J**). We then investigated whether the loss of SOX9+ cells could be caused by cell death in the NCC-*Fgfr2^-/-^* mice. Immunofluorescence for cleaved-caspase showed no cell death in controls or NCC-*Fgfr2^-/-^*mutants at P0, P3, or P5 (**Supplemental Figure 1**). All together these results strongly indicate that the SCX+ cells of the AF connective tissue normally undergo regionally selective differentiation to contribute to intramembranous bone, endochondral bone, and connective tissue in the PFS, and that the persistent AF in NCC-*Fgfr2^-/-^* mice is largely caused by failed differentiation of SCX+ cells into RUNX2+ and SOX9+ skeletogenic cells and retained connective tissue fate.

*Fgfr2* has a well-established role in regulating proliferation and differentiation in the developing sutures. We next investigated how loss of *Fgfr2* impacts proliferation during AF closure using EdU pulse-chase experiments. At E18.5, no statistically significant differences were identified in the percentage of proliferating cells within the AF connective tissue or osteogenic fronts of the paired frontal bones (p>.02) (**Figure 7A-C**). When these regions were combined, a statistically significant difference in proliferation of less than 1% is seen between control and NCC-*Fgfr2^-/-^* mice (6% in total control versus 5.1% in total mutant, p=0.02, n=7) (**Figure 7C**). At P3, EdU pulse-chase shows a significant difference in proliferation of 2.3% between control and mutant mice (9.6% in total controls versus 7.3% in total mutant, p=.004, n=6) (**Figure 7D-F**). Interestingly, this difference can primarily be accounted for by a decrease in the proliferative rate of the NCC-*Fgfr2^-/-^*AF connective tissue cells (6.8% in control versus 4.9% in mutants, p=.001, n=6). Proliferation rates in the bone fronts were not statistically significantly different between mutant and control (**Figure 7F**). Together this indicates that while proliferation within bone front cells appeared normal, a small decrease in the proliferation rate of AF connective tissue could contribute to AF patency in NCC-*Fgfr2^-/-^* mice but does not account for the changes seen in cellular differentiation.

**Figure 7.**
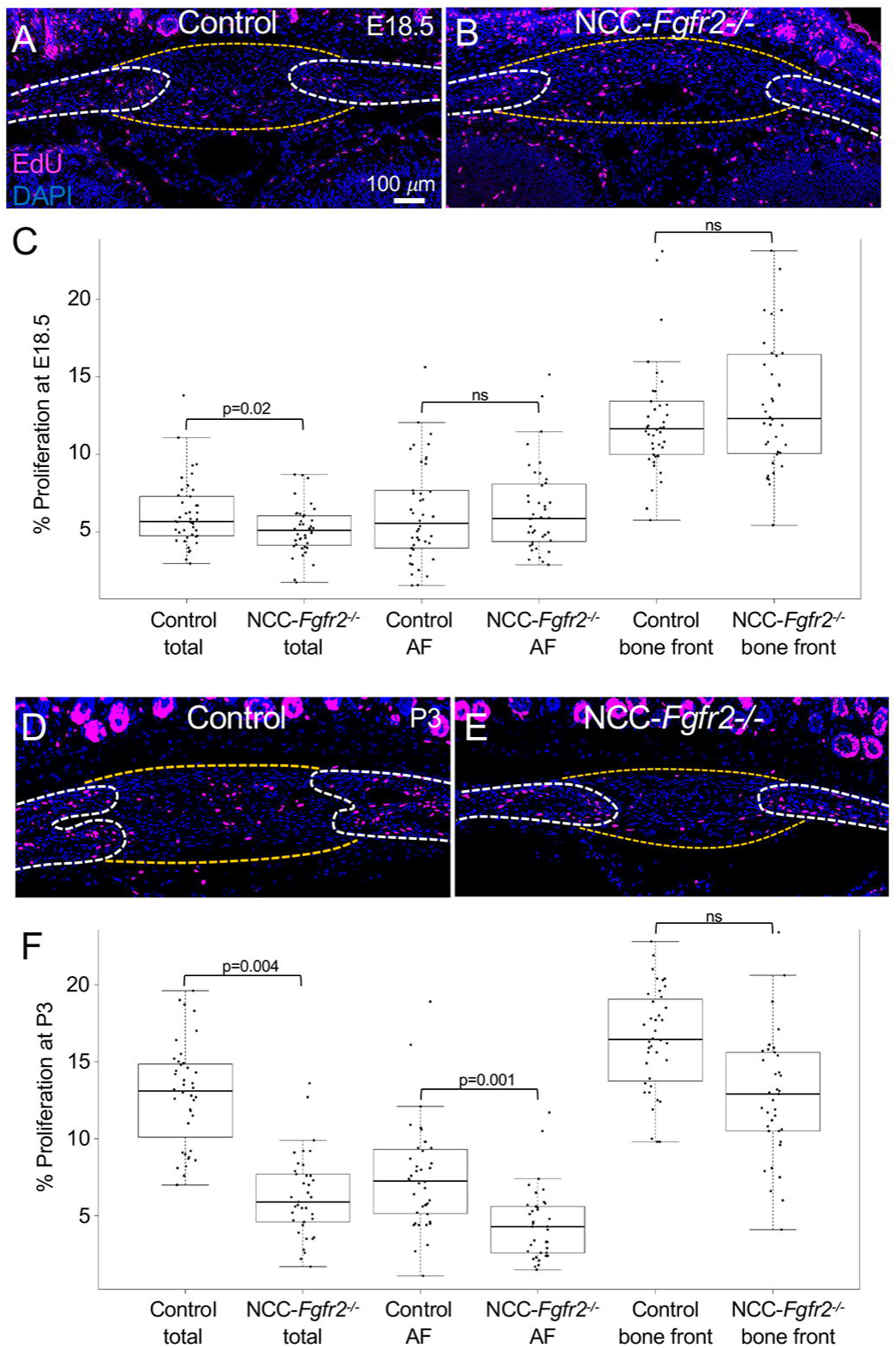
EdU pulse chase identifies a minor decrease in proliferation in the AF of NCC-*Fgfr2^-/-^*mutant mice. (A-C) EdU pulse chase at E18.5 in control (A,C) and NCC-*Fgfr2^-/-^*littermates (B,C) identified a similar average percent of EdU+ (pink) proliferating cells within the osteogenic bone fronts (white dotted lines) and AF (yellow dotted lines). A significant difference in proliferation is only seen when counting both the bone front and AF regions (6% in control vs. 5.1% in mutant, p=.02) (n = 7 littermate pairs). (D-F) At P3, total proliferation was elevated in control samples (9.6% in control vs. 7.3% in mutant, p= .004) with the bulk of this difference occurring in the AF connective tissue (6.8% in control vs. 4.9% in cKO, p= .001) (n = 6 littermate pairs). Statistical significance was calculated using a Student’s t-test assuming unequal variance.

### Cells of the anterior fontanelle are non-autonomously regulated by *Fgfr2*

To determine the fate of AF cells during PFS formation, we traced *Scx-*lineage (Scx^LIN^) and *Sox9*-lineage (Sox9^LIN^) cells at P4 and P7. In *Scx-Cre;TdTomato:Ai9* mice at P4, Scx^LIN^ cells occupy the osteogenic fronts of the frontal bones, as well as the ecto- and endocranial layers of the AF (**Figure 8A**). At P7, the contribution of Scx^LIN^ cells to the ecto- and endocranial layers of the AF has increased, occupying the mid-suture, osteogenic fronts of the frontal bones, and the suprasutural ligament (**Figure 8B**). In *Sox9-CreERT2;TdTomato:Ai9* mice induced with tamoxifen at P2 and P3, Sox9^LIN^ cells are enriched in the endocranial domain of the AF and also contribute to the frontal bones at P4 (**Figure 8C**). At P7, the Sox9^LIN^ cells contribute to the condensing cartilage in the endocranial domain of the PFS (**Figure 8D**). These results show that different cell populations of the AF make direct and region-specific contributions to the PFS.

**Figure 8.**
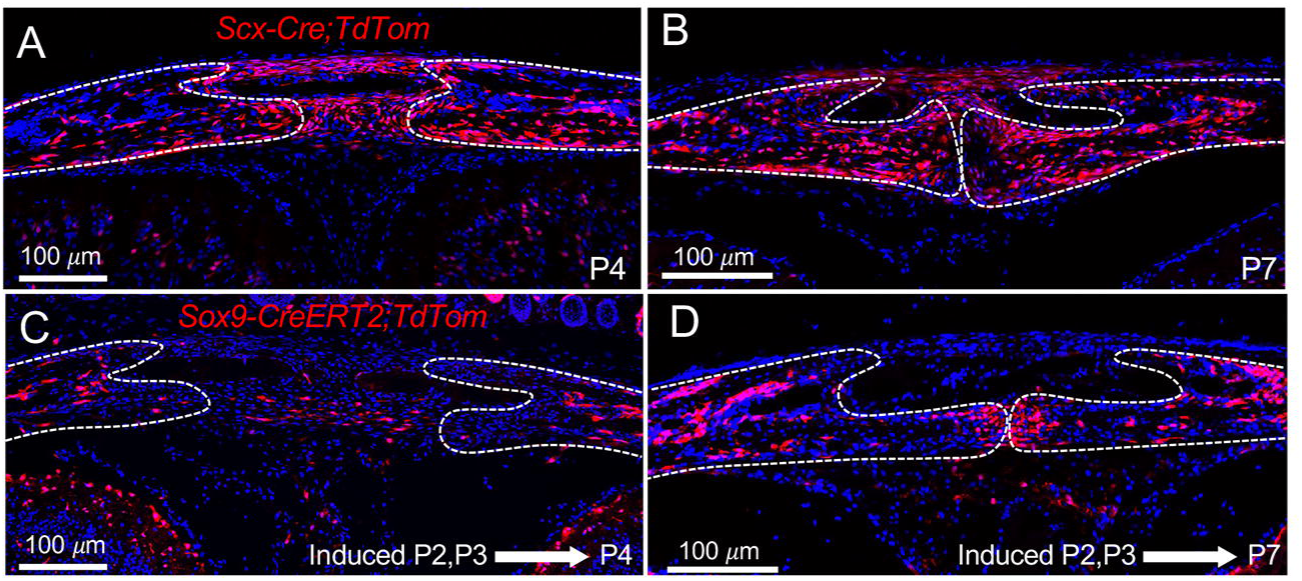
*Scx*-lineage and *Sox9*-lineage directly contribute to the developing PFS. (A) Genetic lineage tracing in *Scx-Cre;TdTomato:Ai9* mice show widespread contribution of Scx^LIN^ cells at P4 and P7 to both the osteogenic fronts of the frontal bones and the AF connective tissue. (B) At P7, when the PFS is established, Scx^LIN^ cells contribute to the ecto- and endocranial bone, as well as the suprasutural ligament. (C) Genetic lineage tracing in *Sox9-CreERT2;TdTomato:Ai9* mice following tamoxifen induction at P2 and P3, identifies Sox9^LIN^ in the osteogenic fronts of the frontal bones and within the endocranial domain of the AF at P4. (D) At P7, the Sox9^LIN^ cells contribute to the cartilage condensation within the endocranial domain. Dotted lines in each panel outline osteogenic fronts. (N = 3 per stage)

We next tested the extent to which *Fgfr2* is required in *Scx* and *Sox9* expressing cells during AF development. Wholemount skeletal staining of *Scx-Cre;Fgfr2^-/-^* mice at P5 and P7 showed that the AF closure is indistinguishable from littermate controls **(Figure 9A-D**). Since feature plots of the scRNA-seq dataset identify cells within the AF1 and AF2 clusters that co-express *Fgfr2* and *Scx* (**Figure 9E**), we conclude that loss of *Fgfr2* in *Scx* + cells is not sufficient to recapitulate the AF and PFS phenotypes seen in NCC-*Fgfr2-/-* mice. Wholemount skeletal staining of *Sox9-CreERT;Fgfr2^-/-^* mice at P5 following tamoxifen induction at E17.5 and 18.5 revealed that AF closure is indistinguishable from littermate controls (**Figure 10A,B**). The same result was found when *Sox9-CreERT; Fgfr2^-/-^* mice were induced at P0 and collected at P7 (**Figure 10C,D**). However, histological examination of the PFS in *Sox9-CreERT; Fgfr2^-/-^* mice at P9 following tamoxifen inductions at P2 and P3 shows loss of the endocranial cartilage (**Figure 10E,F**). Feature plots of the scRNA-seq dataset identify cells within the AF2 and OF2 clusters that co-express *Fgfr2* and *Sox9* (**Figure 10G**), and therefore, while there is some autonomous requirement for *Fgfr2* during endochondral-like fusion of the PFS, it is not required in *Sox9*+ cells for AF closure.

**Figure 9.**
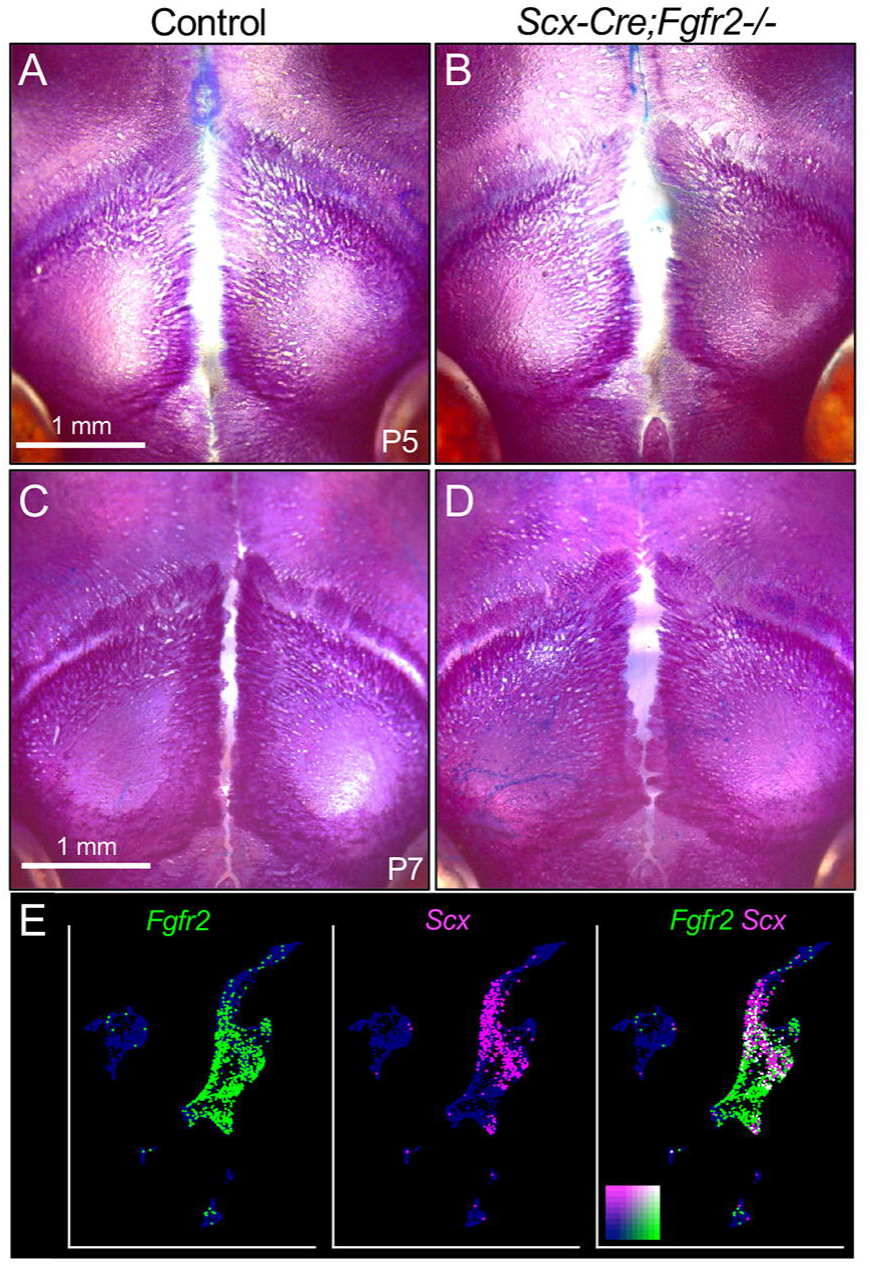
*Fgfr2* is not required in *Scx*+ cells during AF closure. (A-D) Wholemount alizarin red and alcian blue staining at P5 and P7 shows *Scx-Cre;Fgfr2^-/-^* mice resemble littermate controls, showing no difference in the closure of the AF and development of the PFS. (N = 3 littermate pairs per stage) (E) Co-expression analysis in the scRNA-seq data indicates that *Scx*+ cells within the AF1 and AF2 clusters express *Fgfr2* at E18.5. Individual cells are color coded as either *Fgfr2+* only (green), *Scx+* only (pink), *Fgfr2+/Scx+* simultaneously (white) or *Fgfr2-/Scx-* (blue).

**Figure 10.**
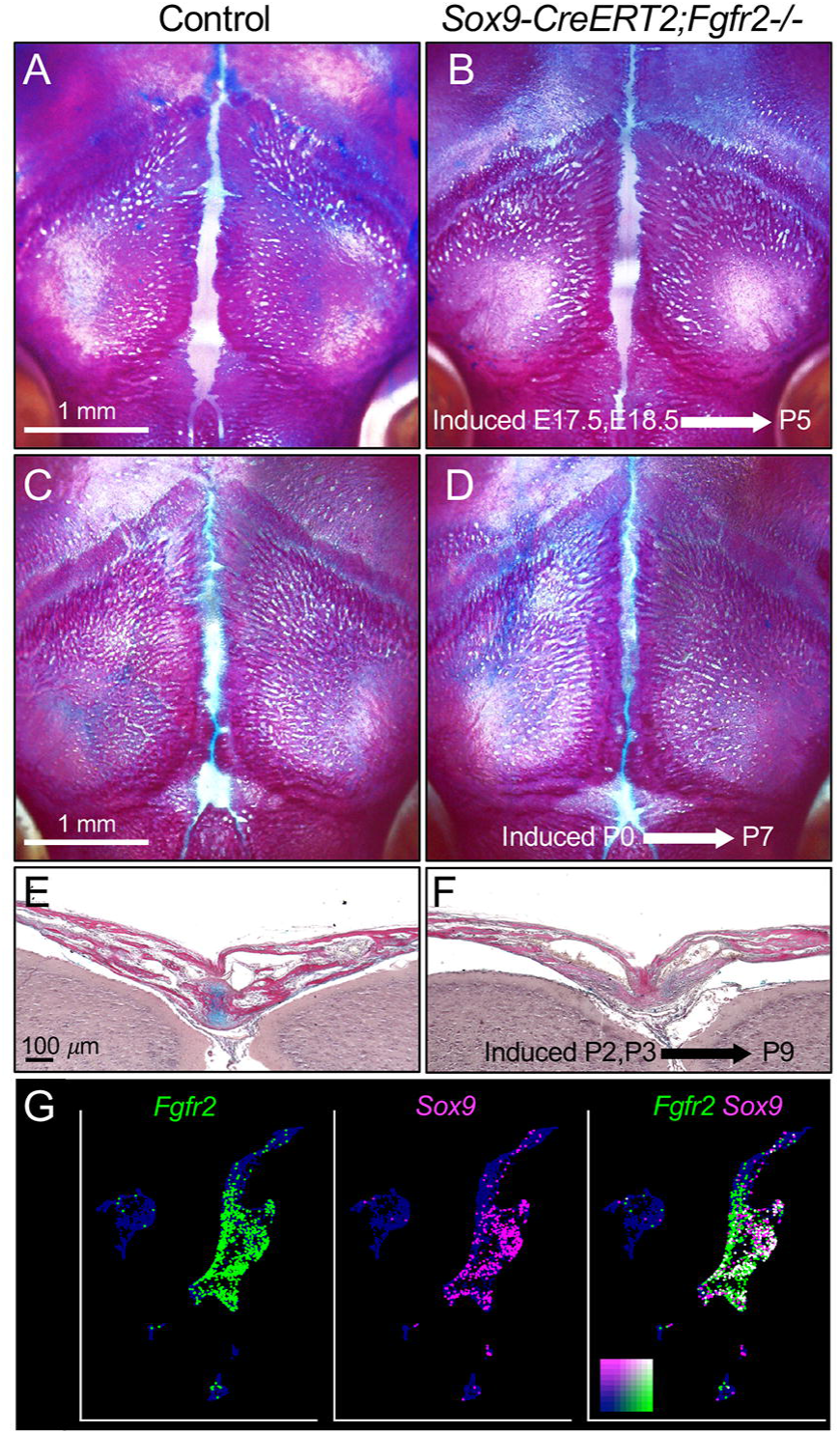
*Fgfr2* is not required in *Sox9*+ cells during AF closure. (A,B) Wholemount alizarin red and alcian blue staining in *Sox9-Cre-ERT2;Fgfr2^-/-^*and littermate controls at P5 following tamoxifen induction at E17.5 and E18.5 shows no difference in AF closure. (C,D) Wholemount alizarin red and alcian blue staining in *Sox9-Cre-ERT2;Fgfr2^-/-^* and littermate controls at P7 following tamoxifen induction at P0 shows no difference in AF closure. N = 3 littermate pairs per stage. (E-F) HBQ staining of the PFS at P9 following tamoxifen induction at P2 and P3 shows loss of cartilage in the endocranial layer in *Sox9-Cre-ERT2;Fgfr2^-/-^*mutants compared to littermate controls (N = 4 littermate pairs). (G) Co-expression analysis in the scRNA-seq data shows *Sox9*+ cells within the AF2 and OF2 clusters express *Fgfr2* at E18.5. Individual cells are color coded in the same manner as Figure 9.

### Fgfr2 regulates expression of WNT pathway members in the anterior fontanelle

To determine the molecular mechanisms through which *Fgfr2* regulates differentiation within the AF, we performed bulk RNA-seq of the control and NCC-*Fgfr2^-/-^*AF at E18.5, when a phenotypic difference is first detected. Heat map analysis of the differentially expressed genes shows that RNA expression profiles of control and NCC-*Fgfr2^-/-^* littermates have significant, genotype specific differences (**Figure 11A**). GO analysis of the genes indicates differential expression of members of the WNT signaling pathway, as well as genes related to focal adhesion, ECM-receptor interaction, and axon guidance (**Figure 11B**). Previous studies have identified a role for WNT signaling in the developing suture, where this pathway works cooperatively with FGF signaling to determine skeletogenic cell fate (Liao et al., 2022, Quarto et al., 2010, Maruyama et al., 2010). Dot plots of WNT pathway members shows that secreted inhibitors *Sfrp4*, *Notum*, and *Wif1*, along with the WNT potentiator *Lgr6* are downregulated in the NCC-*Fgfr2^-/-^* AF (**Figure 11C**). On the other hand, the WNT//J-catenin target gene and WNT signaling potentiator *Lgr5*, *Wnt11*, and the context dependent WNT modulator *Sfrp2* are upregulated in the NCC-*Fgfr2^-/-^* AF (**Figure 11C**). String analysis of the WNT pathway genes found in our sequencing predicts likely interactions between the FGF and WNT pathways are likely (**Figure 11D**).

**Figure 11.**
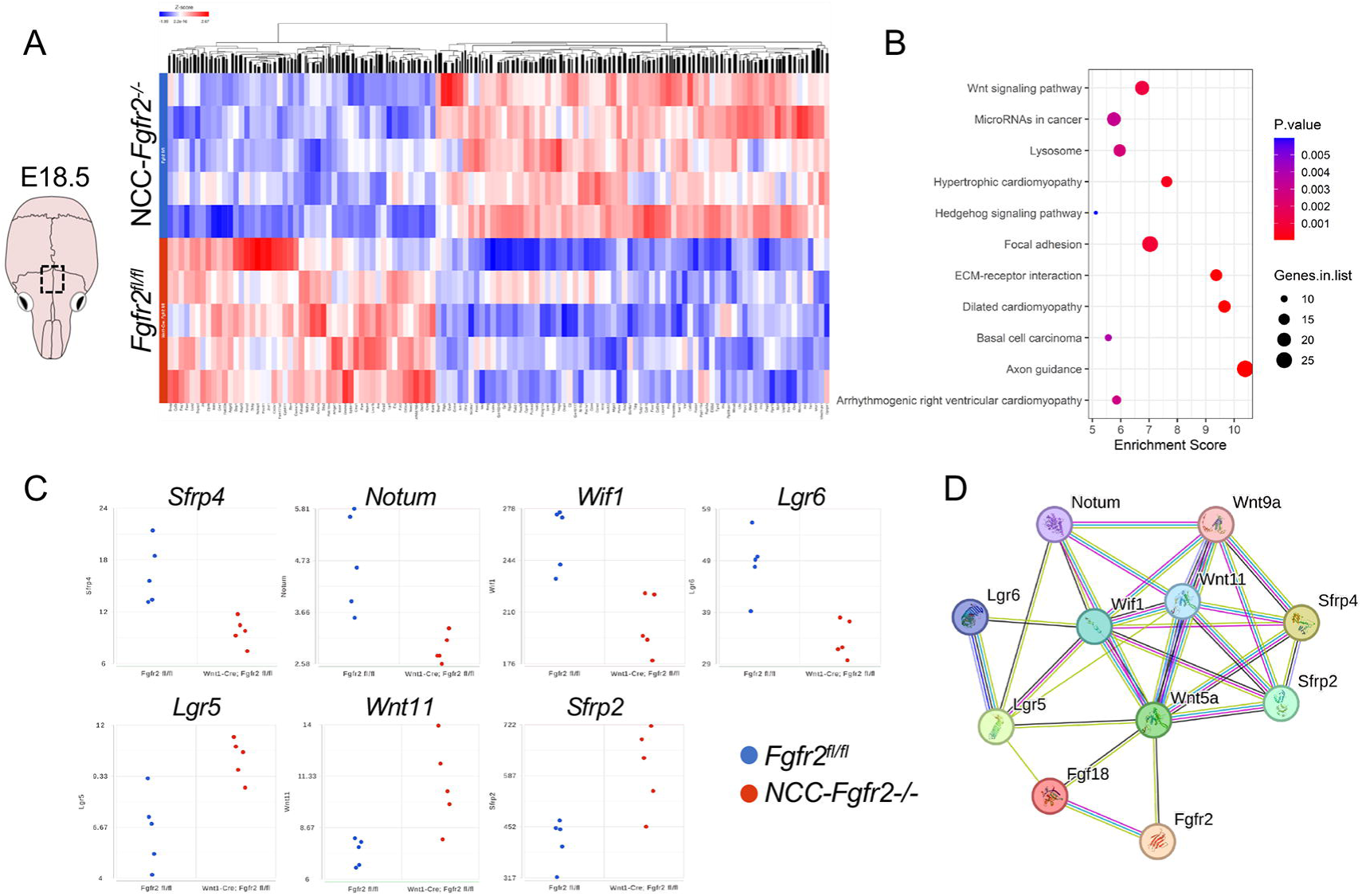
Bulk RNA-seq reveals transcriptional differences in the AF of NCC-*Fgfr2^-/-^* embryos. (A) Diagram of the E18.5 mouse calvarium identifies the region (box) that was analyzed. Heatmap analysis reveals overall differences in gene expression between the AF of NCC-*Fgfr2^-/-^* (top rows) and littermate controls (bottom rows) (n=5 each for control and *NCC-Fgfr2-/-*). (B) GO analysis outlines significant differential regulation of genes associated with focal adhesion, axon guidance, and WNT signaling. (C) Dot plots for differentially expressed genes associated with the WNT pathway. *Sfrp4*, *Notum*, *Wif1, and Lgr6* are downregulated in *NCC-Fgfr2^-/-^* mutants compared to controls, while *Lgr5, Wnt11,* and *Sfrp2* are upregulated in *NCC-Fgfr2^-/-^* mutants compared to controls. (D) String analysis infers interactions between these differentially expressed WNT pathways members with *Fgfr2* and *Fgf18*.

Violin plots of WNT pathways members from the scRNA-seq dataset show that the AF1, AF2, and EM clusters have enriched expression of the WNT receptor *Fzd1*, as well as *Wnt9a*, *Wnt11,* and *Lgr5* (**Figure 12A, A’**). Cellular communication modeling using CellChat predicts that WNT ligands from the EM regulate WNT signaling in AF1 and AF2 (**Figure 12B**). The OF1, OF2, and OB clusters, on the other hand, are enriched for *Wif1* in cells that also express *Fgfr2* (**Figure 12A,A’,C**). WNT-related gene expression changes identified through RNA-seq were validated using RNAScope *in situ* hybridization at E18.5. *Lgr5* expression, which is enriched in the endocranial domain of the control AF, is expanded in the NCC-*Fgfr2^-/-^* AF (**Figure 12D,E**). *Lgr6* expression, which is expressed in both the endo- and ectocranial domains of the control AF, is reduced within the endocranial domain of the NCC-*Fgfr2^-/-^* AF (**Figure 12F,G**). *Wif1* expression is localized to cells within the osteogenic fronts in the control and greatly reduced in NCC-*Fgfr2^-/-^*embryos (**Figure 12H,I**). Together these results indicate that *Fgfr2*-dependent activation of *Wif1* expression in cells of the osteogenic front non-autonomously modulate WNT signaling in *Lgr5* and *Lgr6* expressing cells of the AF. LGR5 is a known stem cell marker in multiple epithelial tissues, and in this context likely represents connective tissue progenitors (Leung et al., 2020, Ng et al., 2014, Shi et al., 2012).

**Figure 12.**
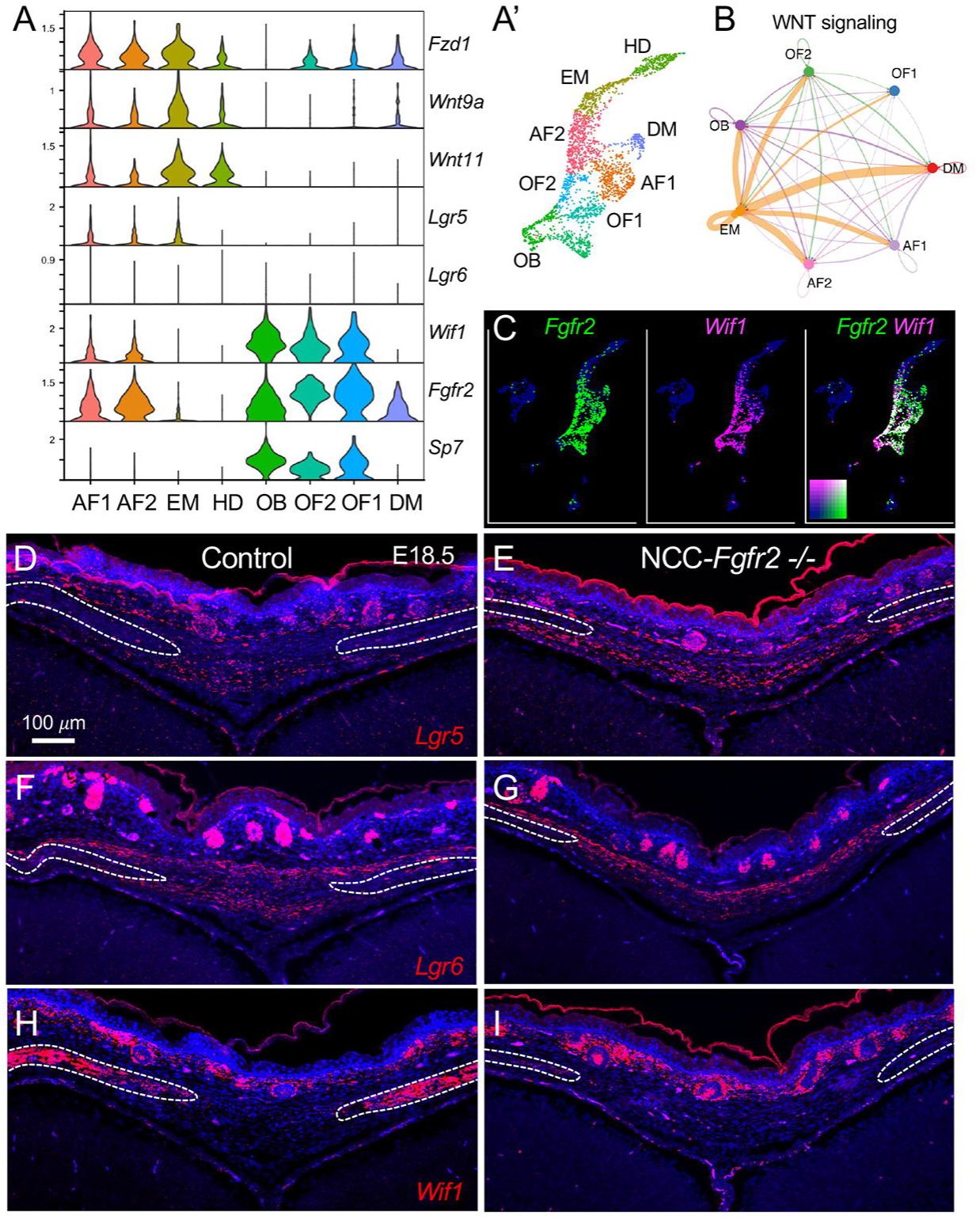
Differentially expressed WNT pathway members correspond to distinct populations within the AF. (A,A’) Violin plots of WNT pathway members show their cluster-specific enrichment in the developing AF at E18.5, with corresponding UMAP shown in panel A’. Expression of genes encoding *Fzd1*, WNT ligands *Wnt9a* and *Wnt11*, and the WNT signaling target *Lgr5* are enriched in the AF1, AF2, and EM clusters, while expression of the WNT inhibitor *Wif1* is enriched in osteogenic populations OF1, OF2, and OB. (B) Cell communication analysis using CellChat predicts EM is a critical source of WNT ligand for AF1 and AF2. (C) Co-expression analysis of *Fgfr2* and *Wif1* shows significant overlap in OF1 and OF2 cluster. Individual cells are color coded in the same manner as Figure 9. (D,E) RNAscope *in situ* hybridization in the AF at E18.5 identifies low level expression of *Lgr5* throughout both ecto- and endocranial domains in the control. In the *NCC-Fgfr2^-/-^*mutant AF, the domain of *Lgr5* is expanded laterally towards the osteogenic fronts. (F,G) While *Lgr6* is expressed throughout the ecto- and endocranial domains of the AF in control, expression within the endocranial domain is lost in the *NCC-Fgfr2^-/-^* mutant. (H,I) In control, *Wif1* is expressed throughout the osteogenic fronts. In the *NCC-Fgfr2^-/-^*mutant, *Wif1* expression is greatly decreased. N = 3 littermate pairs.

### Modulation of WNT signaling impacts anterior fontanelle closure

To test the impact of increased WNT//J-catenin signaling in *Scx* + cells of the AF, we crossed mice harboring the *Scx-Cre* driver to mice with a stabilizing mutation in exon 3 of/J-*catenin* (*Ctnnb1^lox(Ex3)^*) (Harada et al., 1999). Stabilization of the β-CATENIN protein blocks its degradation through the destruction complex and mimics conditions of constitutively active WNT signaling. Wholemount skeletal staining of *Scx-Cre; Ctnnb1^lox(Ex3)/+^* (*Scx-Cre;Ctnnb1^GOF^*) mice at E18.5 shows that the AF is larger compared to littermate controls (**Figure 13A,B**). At P5, when the frontal bones of control mice have approximated to form the PFS, the AF remains patent and the PFS fails to form in *Scx-Cre;Ctnnb1^GOF^* mice (**Figure 13C,D**). The observation that increased WNT//J-catenin signaling in *Scx*+ cells recapitulates the phenotype of NCC-*Fgfr2^-/-^*mice supports a mechanistic connection between FGF and WNT signaling during AF closure and PFS formation.

**Figure 13.**
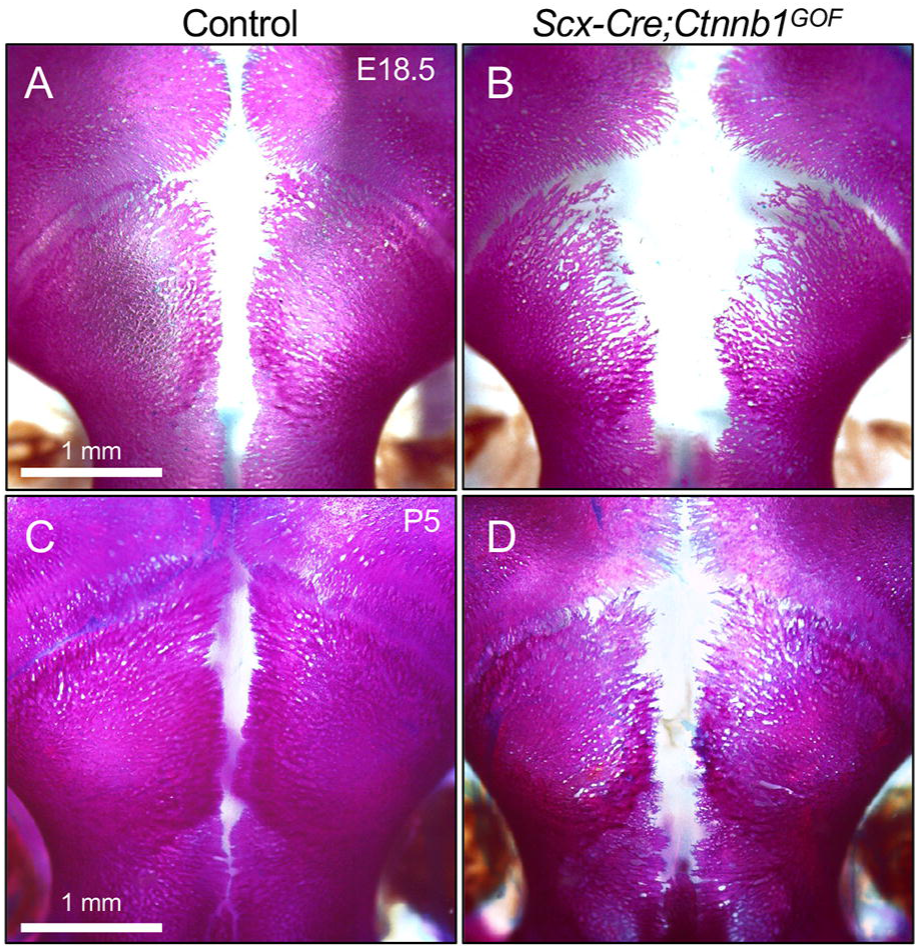
Constitutive activation of WNT//J-catenin signaling in *Scx*+ cell blocks AF closure. (A-D) Wholemount alizarin red and alcian blue staining at E18.5 and P5 show *Scx-Cre;Ctnnb1^GOF^* mice exhibit a wide-open AF that fails to close compared to littermate controls. N = 3 littermate pairs per stage.

To test the idea that elevated WNT signaling in NCC-*Fgfr2^-/-^*mice underlies the AF closure defect, we attempted a rescue through WNT signaling inhibition. To inhibit WNT signaling, we used a cocktail of mouse WIF1 recombinant protein and Vantictumab, a humanized monoclonal antibody that inhibits WNT pathway signaling through competitive binding of FZD receptors. At P3, NCC-*Fgfr2^-/-^*pups and littermate controls were treated with a single injection of the WNT inhibitor cocktail just under the scalp at the location of the AF. Samples were collected at P7 and processed for wholemount skeletal staining. Following treatment, AF closure was largely restored in NCC-*Fgfr2^-/-^* mutants, indicating that elevated WNT signaling underlies the phenotype and is a critical driver of its mechanism (**Figure 14A,B**).

**Figure 14.**
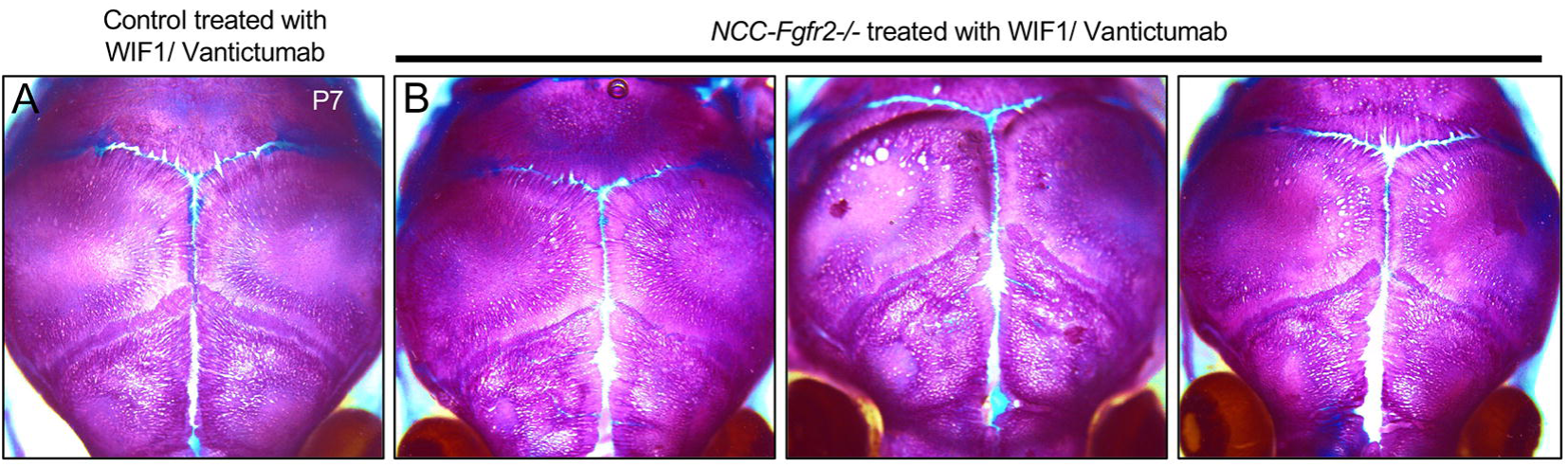
Local inhibition of WNT signaling partially restores AF closure in NCC-*Fgfr2^-/-^* mice. A WNT inhibitor cocktail composed of recombinant mouse WIF1 protein and Vantictumab was administered at P3 through a single injection under the scalp in the area above the AF. (A) Wholemount alizarin red and alcian blue staining of treated controls collected at P7 show that treatment has no negative effects on calvarial bone formation and AF closure. (B) Wholemount skeletal stains of three treated *NCC-Fgfr2^-/-^*littermates show that AF closure is partially restored. N = 7 littermate pairs.

## Discussion

This study characterizes a novel role for FGF signaling in calvarial development. Using a combination of mouse genetics and single cell transcriptomics, we demonstrate that *Fgfr2* is necessary for closure of the anterior fontanelle and development of the posterior frontal suture it prefigures. We find that cells of the anterior fontanelle, which are marked by *Scx* and differentially express *Sox9*, are regionally organized into ecto- and endocranial domains that give rise to distinct lineages within the posterior frontal suture. We provide evidence that FGFR2 signaling in the osteogenic fronts of the frontal bones, through induction of the expression of *Wif1*, induces local inhibition of WNT signaling in the anterior fontanelle (**Figure 15**). Within the zone of WNT inhibition, SCX+ cells in the ectocranial domain are recruited to form bone, while SCX+/SOX9+ cells in the endocranial domain are recruited to form cartilage. Ectocranial SCX+ cells that remain outside the domain of WNT inhibition give rise to suprasutural ligament. Upon loss of FGFR2 signaling, *Wif1* expression is lost, WNT signaling remains, and cells of the anterior fontanelle retain a connective tissue-like fate instead of forming the posterior frontal suture.

**Figure 15.**
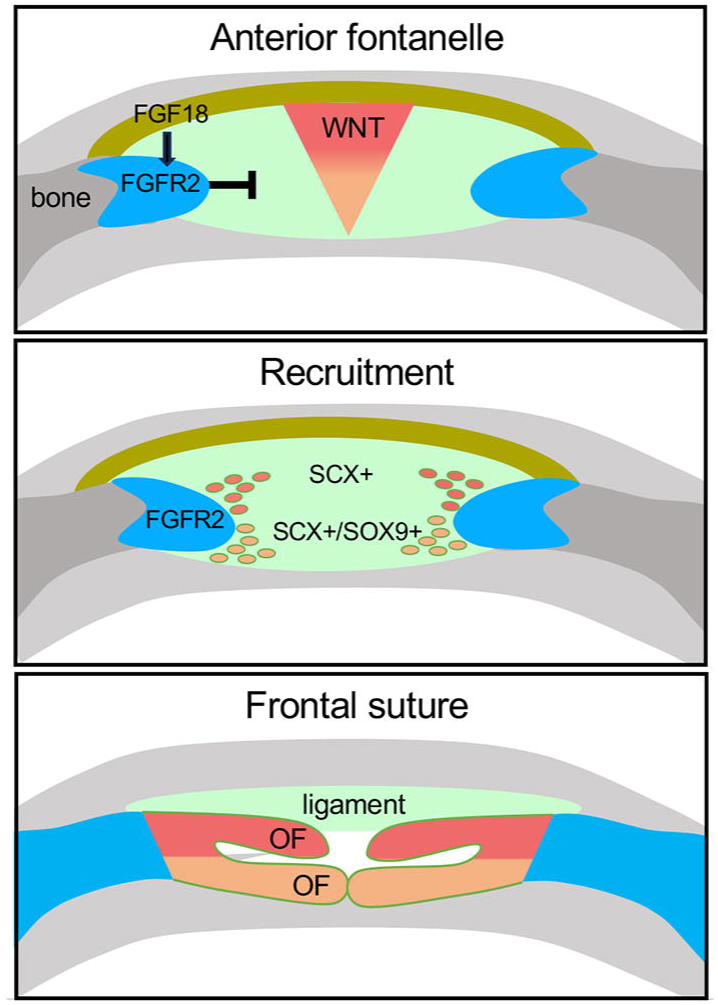
Recruitment model for frontal suture formation within the anterior fontanelle. A gradient of WNT signaling maintains cells of the anterior fontanelle in an undifferentiated state. As FGFR2+ cells of the osteogenic front advance, they secret WIF1 to locally inhibit WNT signaling in the anterior fontanelle. Downregulation of WNT signaling induces SCX+ cells in the ectocranial domain to form intramembranous bone, and SCX+/SOX9+ cells in the endocranial domain to form cartilage. Recruited SCX+ cells form the frontal suture, while SCX+ cells that remain outside the domain of WNT inhibition give rise to suprasutural ligament. Upon loss of FGFR2 signaling, WNT signaling remains high and AF cells fail to form the frontal suture joint.

Our results reveal that FGF and WNT pathways interact to coordinate frontal bone advancement with suture joint formation. We show that differential expression of FGF and WNT pathway members in cells of the ectocranial mesenchyme, osteogenic fronts, and anterior fontanelle establish a signaling center that promotes unique lineage potential along the ectocranial-endocranial axis. Ectocranial mesenchymal cells express FGF18 and WNT ligands, suggesting these cells are a critical source of secreted signals that act on *Fgfr2*+ cells in the osteogenic fronts and *Fgfr2*+ and *Fzd1*+ cells in the anterior fontanelle. FGF18 likely acts to promote osteoblast cell fate in *Fgfr2*+ cells, which is consistent with prior studies showing that loss of *Fgf18* leads to wide-open fontanelles and ectopic FGF18 accelerates approximation of the bone fronts in the coronal suture (Ohbayashi et al., 2002, Sahar et al., 2005, Nagayama et al., 2013). WNT, on the other hand, likely acts to maintain *Fzd1*+ cells in the anterior fontanelle in an undifferentiated state. This is supported by multiple studies showing that, in the posterior frontal suture, canonical WNT signaling is dynamic and sharply down-regulated immediately preceding cartilage formation in the endocranial domain while continuous activation of WNT leads to a wide-open frontal suture and failure of cartilage formation (Behr et al., 2010, Behr et al., 2011, Behr et al., 2013). We propose that *Fgfr2*+ cells at the osteogenic fronts downregulate WNT in the anterior fontanelle to promote chondrogenic fate through their expression of the secreted WNT inhibitor WIF1. That WNT promotes connective tissue over cartilage fate is also supported by studies in the developing limb (ten Berge et al., 2008).

The calvaria develops from a layer of head mesenchyme that is derived from mesoderm or neural crest, and regionally patterned into osteogenic and non-osteogenic domains. Mesenchyme that initially resides in the supraorbital ridge just above the eyes, called the supraorbital mesenchyme (SOM), generates the ossification centers for the frontal and parietal bones (Ishii et al., 2015, Deckelbaum et al., 2012, Jiang et al., 2002, Yoshida et al., 2008). In contrast, mesenchyme that lies apical to the SOM, called the early migrating mesenchyme (EMM), remains non-osteogenic through embryonic stages of calvarial development (Cesario et al., 2018, Roybal et al., 2010). Growth of the frontal and parietal bone rudiments towards the vertex of the head in embryogenesis is fueled by the SOM through intrinsic growth rather than recruitment of adjacent EMM mesenchyme (Yoshida et al., 2008). Instead, the EMM contributes to non-osteogenic connective tissue that occupies the sutures and fontanelles (Cesario et al., 2018). Our findings elaborate on this model by showing that, in the postnatal period, non-osteogenic SCX+ cells within the AF directly contribute to bone, cartilage, and ligament within the PFS. This suggests that formation of the PFS within the anterior fontanelle is developmentally distinct. Our results suggest that, as the frontal bones approximate, they recruit SCX+ cells of the AF to initiate frontal suture joint formation, which is characterized by the formation of a secondary osteogenic front within the endocranial layer that subsequently undergoes endochondral fusion. This may be a key factor that distinguishes the bilayered, fusing murine PFS from the overlapping coronal suture or abutting sagittal suture, which do not fuse during normal mouse development. Importantly, SCX+ cells in the AF appear to be distinct from those we previously identified in the non-fusing murine coronal suture. In the coronal suture, SCX+ cells are highly restricted to the overlapping edges of the frontal and parietal bones in the ectocranial domain and appear to remain non-osteogenic (Farmer et al., 2021).

Overall, the results presented in this study offer further insight into regional differences in the development of sutures that are prefigured by fontanelles. It also suggests a possible etiology for the co-occurrence of premature suture fusion and along with persistent AF patency observed in craniosynostosis syndromes such as those caused by *FGFR2* GOF. While it is not yet clear how FGFR2 GOF impacts the mechanism for AF closure we described here, a previous study found that *Lgr5* is upregulated in the developing frontal suture of *Twist+/-* embryos, a model for Saethre-Chotzen syndrome that present with persistent AF and premature coronal suture fusion (Holmes et al., 2020).

## Materials and Methods Mice

To conditionally knockout *Fgfr2* in NCC derived cells, *Fgfr2^flx/flx^*(JAX Stock No. 007569) mice were crossed with the *Wnt1-Cre* driver (JAX Stock No. 003829), the *Sox9-CreERT2* driver (JAX 035092) (Akiyama et al., 2005), or the *Scx-Cre* driver which was generously provided by Dr. Ronen Schweitzer. The *Fgfr2^flx/flx^* and *Wnt1-Cre* lines were previously described (Lewis et al., 2013, Yu et al., 2003). *Scx-GFP* was used to mark dense connective tissue fibroblasts (Pryce et al., 2007). *Scx-Cre,* and *Sox9-Cre-ERT2* lines were described previously (Xu et al., 2015, Sugimoto et al., 2013). The *Ai9* allele (JAX Stock No. 007909) was used as a lineage marker for those tissues targeted by *Scx-Cre,* and *Sox9-Cre-ERT2* (Madisen et al., 2010). For *Sox9-Cre-ERT2* mice, Tamoxifen (Sigma; dissolved in peanut oil ON @40°C) was delivered via oral gavage to either timed pregnant females (for embryonic inductions; 1mg/induction/dam) or neonatal mice (for postnatal inductions; 0.1mg/induction/pup). Finally, constitutive activation of/J*-catenin* in fibroblasts was driven by crossing *Ctnnb^lox(Ex3)^* mice (Harada et al., 1999) to mice carrying the *Scx-Cre* driver. Embryonic samples were collected from timed pregnant females. Postnatal samples were staged according to the date of birth. All experimental protocols were approved by the USC’s Institutional Animal Care and Use Committee.

### Skeletal preparation

Mice processed for wholemount imaging were collected as above and following PBS rinse were fixed in 95% ethanol for a minimum of 4 days. Fixed samples were stained for cartilage using 0.15mg/mL Alcian Blue (Sigma-Aldrich CAT. A5268) in 80% ethanol and 20% glacial acetic acid. Samples were then de-stained in 95% ethanol for a minimum of two days. Tissue clearing was performed using KOH between 0.5-1% w/v for between 2-8 days depending on size of sample. Calcified tissue was then discriminated using 0.02 mg/mL Alizarin Red S (Sigma-Aldrich CAT. A5533) in 0.5-1% KOH. Destaining and further clearing was performed as needed in 0.5-1% KOH. Samples were equilibrated, stored, and imaged in 75% glycerol. Measurements of the AF area using ImageJ and each individual area measurement was normalized by the length of the PFS from the jugum limitans to the coronal suture. These were then averaged within each group and the average AF area of control and NCC-*Fgfr2-/-* mice was compared by age using an unpaired, two-tailed Student’s t test assuming unequal variance.

### Micro-computed tomography (μCT)

All μCT scans were performed by the USC Molecular Imaging Center using a μCT50 (Scanco Medical). Samples were rotated 360° and X-ray settings were standardized to 90 kV and 155 µA with an exposure time of 0.5 seconds per frame to yield a nominal resolution of 20 μm. A 0.5-mm-thick aluminum filter was employed to minimize beam-hardening artifacts. Morphometric analysis was performed using the Amira 6.2 and VG Studio MAX 3.0 software packages. Isosurface renderings with equal threshold were measured using the 3D measuring tool. All jaw measurements and landmarks were measured as previously described (Guerreiro et al., 2013). These experiments were performed on at least 3 biological replicates, which we defined as 3 same-sex littermate pairs (control and mutant) derived from 3 different litters. Statistical significance was determined using unpaired two-tailed Student’s t-tests.

### Histology

Samples used for histology were processed as follows; animals were humanely sacrificed and decapitated. Heads were then fully or partially skinned, rinsed in PBS, and fixed in 10% neutral buffered formalin (NBF) at room temperature for between 24-36 hours depending on stage. Following fixation, samples were decalcified in 10% EDTA (pH 7.4) at 4°C for 3-7 days, until calvaria began to concave. Samples were then dehydrated in serial ethanol washes of increasing concentration, equilibrated in Citrus Clearing Solvent (VWR CAT.72060-044), and embedded following 3 washes in pure paraffin wax undergone at 65°C with vacuum pressure. Embedded samples were then cut into 8 µm sections. Sections were stained using the Hall-Brunt quadruple stain (Hall, 1986).

### Immunofluorescent analysis

Samples for immunofluorescent staining were harvested and fixed in 4% PFA at 4°C for between 30-60 mins depending on stage. Decalcification of bone was performed via incubation in 10% EDTA (pH 7.4) at 4°C for between 3-7 days until calvaria began to concave, Subsequently, samples were equilibrated sequentially in 15% and 30% sucrose/PBS at 4°C until they sank, then embedded in optimal cutting temperature (O.C.T.) compound (Electron Microscopy Sciences). Cryosectioning was performed at a section thickness of 8μm onto Superfrost+ slides. Sections were washed three times with PBS, permeabilized with 0.1% TritonX-100 in PBS, then blocked for one hour at RT in 10% serum (either donkey or goat from Sigma). Slides were incubated in primary antibody overnight at 4°C (**Supplemental Table 1**), then washed in PBS and incubated in Alexa Fluor 568 secondary antibody (ThermoFisher Scientific A-11036) at a concentration of 1:200 for one hour at RT. Finally, slides were washed in PBS and mounted in Vectashield with DAPI (Vector Labs). Slides used for lineage tracing were washed three times in PBS and counterstained with DAPI as described above. Slides were imaged using either a Leica TCS SP5/8 or Stellaris 5 confocal system. These experiments were performed on 3 biological replicates, which we defined as 3 littermate pairs (control and mutant) derived from 2 different litters.

### Single cell transcriptomics

Data for scRNA-seq analysis was obtained from FaceBase (Accession #1-4TT6) (Holmes et al., 2020). Seurat 3 R-Package was used to explore QC metrics and filter for high-quality cells (Stuart et al., 2019). The filtered dataset representing 3366 cells (median of 3750 genes per cell) was normalized (log normalized), scaled (linear transformation), and clustered into 16 clusters (dims 1:13, resolution = 0.7) using unsupervised graph-based clustering based on principal component analysis scores. The identities of the clusters were resolved using previously reported markers. Pseudotime analysis was conducted using Monocle 3 (Cao et al., 2019). Cell-cell communication predictions were made using CellChat (Jin et al., 2021).

### *In situ* hybridization

Samples for *in situ* hybridization were collected and processed in the same manner as histological samples, described above. Transcripts were detected using RNAscope Fluorescent Multiplex Assay v2 (ACD) as per manufacturer’s instructions. Briefly, slides with paraffin sections were baked in an embedding oven for 60 mins. at 65°C. Slides were then deparaffinized using xylenes, dehydrated in 100% EtOH, and allowed to completely air dry. Endogenous peroxidase activity was quenched using hydrogen peroxide provided within the kit and slides were washed in deionized water. Antigen retrieval was performed using the provided Target Retrieval Reagent for 15 mins. in an Oster steamer heated to 99°C. Slides were then dehydrated in 100% EtOH and allowed to completely air dry. The slides were treated with ACD Protease plus in a humidified slide chamber for 15 mins. at 40°C. Probe hybridization was also performed in a 40°C humidified slide chamber for 2 hours (**Supplemental Table 1**). Slides were then stored in 5x SSC solution overnight. Amplification steps were performed as prescribed by manufacturer and signal development was done using Opal 570 (Akoya Biosciences FP1488001KT) Opal 620 (Akoya FP1495001KT) or Opal 690 (Akoya FP1497001KT) diluted 1:1000 in the ACD provided TSA buffer. Slides were then counterstained using Vectashield mounting medium with DAPI and imaged using the Leica TCS SP5/8 or Stellaris 5 confocal system. These experiments were performed on 3 biological replicates, which we defined as 3 littermate pairs (control and mutant) derived from 2 different litters.

### EdU Proliferation Assay

Cell proliferation was assayed using an *in vivo* EdU Click kit according to manufacturer instructions (Sigma BCK647-IV-IM-M). Briefly, E18.5 samples were obtained by intraperitoneally injecting timed pregnant females with 1.5 mg EdU dissolved in PBS, while P3 mice received subcutaneous injections of 0.1mg EdU dissolved in PBS. In both cases, a chase period of four hours was observed before sample collection. Samples were cryo-embedded and sectioned at 8μm as described above. Detection of EdU took place in the dark, at room temperature for 30 mins. Slides were then counterstained and mounted using Vectashield mounting media with DAPI. For analysis, littermate control and NCC-*Fgfr2-/-* pairs were selected, and a minimum of 6 different sections from each sample were imaged and uploaded to ImageJ. Each image was quantified for total AF cells via individual nuclei, and the number of proliferating cells was counted. Statistical analysis was performed on total proliferation in mutant vs control, proliferation of AF fibrous connective tissue, and proliferation in bone front cells. Statistical significance was determined using unpaired two-tailed Student’s t-tests assuming unequal variance.

### RNA Isolation and Gene Expression Analysis

The region of the anterior fontanelle was excised from E18.5 embryos (including both suture connective tissue and bone but excluding skin and brain) and added to 300µL of DNA/RNA Shield (Zymo). RNA was isolated using the *Quick*-RNA Miniprep Plus Kit (Zymo) according to manufacturer’s instructions with the addition of an optional DNase digestion and a 1 min dounce homogenization step following proteinase-K digestion. Samples were eluted from the column using 80 µL of DNase/RNase-free water and then analyzed via nano-drop and to check concentration and purity. Library preparation and sequencing was performed at the UCLA TCBG facility using the Kapa Stranded Kit (Roche). Sequencing was performed using Hiseq3000 at 1x50 read length and 30 million reads per sample. Differential gene expression analysis was performed using Partek Flow.

### WNT Inhibitor Cocktail Treatment

Stock Vantictumab (Fisher MA5-42006) and recombinant WIF1 protein (R&D 135-WF-050) were diluted in sterile water to a final concentration of 1.1 mg/mL and 0.5 mg/mL respectively. At P3, pups were treated with a single dose of the inhibitor cocktail (a total of 2ug Vantictumab and 2ug Wif1) using a pre-filled 29G insulin syringe. For the treatment, pups were placed on a nitrile glove on ice for 30-45 seconds to slow movement, and the WNT inhibitor cocktail was injected subcutaneously just under the scalp above the area of the AF. Volume of liquid per injection did not exceed 20uL. Following treatment, pups were placed on a heating pad and observed for several minutes to ensure recovery from the procedure before returning to their dams.

## Acknowledgements

This work was supported by the National Institute of Health [#R01DE025222 to A.E.M, # F31 DE032259 to A.N, and #T90DE021982 in support of L.B.].

## References

Akiyama, H., Kim, J. E., Nakashima, K., Balmes, G., Iwai, N., Deng, J. M., Zhang, Z., Martin, J. F., Behringer, R. R., Nakamura, T. & De Crombrugghe, B. 2005. Osteo-chondroprogenitor cells are derived from Sox9 expressing precursors. Proc Natl Acad Sci U S A, 102, 14665–70.

Azoury, S. C., Reddy, S., Shukla, V. & Deng, C.-X. 2017. Fibroblast Growth Factor Receptor 2 (FGFR2) Mutation Related Syndromic Craniosynostosis. International journal of biological sciences, 13, 1479–1488.

Behr, B., Longaker, M. T. & Quarto, N. 2010. Differential activation of canonical Wnt signaling determines cranial sutures fate: a novel mechanism for sagittal suture craniosynostosis. Dev Biol, 344, 922–40.

Behr, B., Longaker, M. T. & Quarto, N. 2011. Craniosynostosis of coronal suture in twist1 mice occurs through endochondral ossification recapitulating the physiological closure of posterior frontal suture. Front Physiol, 2, 37.

Behr, B., Longaker, M. T. & Quarto, N. 2013. Absence of Endochondral Ossification and Craniosynostosis in Posterior Frontal Cranial Sutures of Axin2−/− Mice. Plos One, 8, e70240.

Bradley, J. P., Levine, J. P., Roth, D. A., Mccarthy, J. G. & Longaker, M. T. 1996. Studies in cranial suture biology: Iv. Temporal sequence of posterior frontal cranial suture fusion in the mouse. Plast Reconstr Surg, 98, 1039–45.

Cao, J., Spielmann, M., Qiu, X., Huang, X., Ibrahim, D. M., Hill, A. J., Zhang, F., Mundlos, S., Christiansen, L., Steemers, F. J., Trapnell, C. & Shendure, J. 2019. The single-cell transcriptional landscape of mammalian organogenesis. Nature, 566, 496–502.

Cesario, J. M., Landin Malt, A., Chung, J. U., Khairallah, M. P., Dasgupta, K., Asam, K., Deacon, L. J., Choi, V., Almaidhan, A. A., Darwiche, N. A., Kim, J., Johnson, R. L. & Jeong, J. 2018. Anti-osteogenic function of a Lim-homeodomain transcription factor LMX1b is essential to early patterning of the calvaria. Dev Biol, 443, 103–116.

Deckelbaum, R. A., Holmes, G., Zhao, Z., Tong, C., Basilico, C. & Loomis, C. A. 2012. Regulation of cranial morphogenesis and cell fate at the neural crest-mesoderm boundary by engrailed 1. Development, 139, 1346–58.

Farmer, D. T., Mlcochova, H., Zhou, Y., Koelling, N., Wang, G., Ashley, N., Bugacov, H., Chen, H. J., Parvez, R., Tseng, K. C., Merrill, A. E., Maxson, R. E., Jr., Wilkie, A. O. M., Crump, J. G. & Twigg, S. R. F. 2021. The developing mouse coronal suture at single-cell resolution. Nat Commun, 12, 4797.

Guerreiro, F. D. S., Diniz, P., Carvalho, P. E. G., Ferreira, E. C., Avancini, S. R. P. & Ferreira-Santos, R. I. 2013. Effects of masticatory hypofunction on mandibular morphology, mineral density and basal bone area. Brazilian Journal of Oral Sciences, 12, 205–211.

Hajihosseini, M. K., Wilson, S., De Moerlooze, L. & Dickson, C. 2001. A splicing switch and gain-of-function mutation in FgfR2-IIIc hemizygotes causes Apert/Pfeiffer-syndrome-like phenotypes. Proc Natl Acad Sci U S A, 98, 3855–60.

Hall, B. K. 1986. The role of movement and tissue interactions in the development and growth of bone and secondary cartilage in the clavicle of the embryonic chick. J Embryol Exp Morphol, 93, 133–52.

Harada, N., Tamai, Y., Ishikawa, T., Sauer, B., Takaku, K., Oshima, M. & Taketo, M. M. 1999. Intestinal polyposis in mice with a dominant stable mutation of the beta-catenin gene. Embo j, 18, 5931–42.

Holmes, G., Gonzalez-Reiche, A. S., Lu, N., Zhou, X., Rivera, J., Kriti, D., Sebra, R., Williams, A. A., Donovan, M. J., Potter, S. S., Pinto, D., Zhang, B., Van Bakel, H. & Jabs, E. W. 2020. Integrated Transcriptome and Network Analysis Reveals Spatiotemporal Dynamics of Calvarial Suturogenesis. Cell Rep, 32, 107871.

Holmes, G., Rothschild, G., Roy, U. B., Deng, C.-X., Mansukhani, A. & Basilico, C. 2009. Early onset of craniosynostosis in an Apert mouse model reveals critical features of this pathology. Developmental Biology, 328, 273–284.

Ishii, M., Sun, J., Ting, M. C. & Maxson, R. E. 2015. The Development of the Calvarial Bones and Sutures and the Pathophysiology of Craniosynostosis. Curr Top Dev Biol, 115, 131–56.

Jiang, X., Iseki, S., Maxson, R. E., Sucov, H. M. & Morriss-Kay, G. M. 2002. Tissue Origins and Interactions in the Mammalian Skull Vault. Developmental Biology, 241, 106–116.

Jin, S., Guerrero-Juarez, C. F., Zhang, L., Chang, I., Ramos, R., Kuan, C.-H., Myung, P., Plikus, M. V. & Nie, Q. 2021. Inference and analysis of cell-cell communication using CellChat. Nature Communications, 12, 1088.

Kague, E., Roy, P., Asselin, G., Hu, G., Simonet, J., Stanley, A., Albertson, C. & Fisher, S. 2016. Osterix/Sp7 limits cranial bone initiation sites and is required for formation of sutures. Dev Biol, 413, 160–72.

Leung, C., Murad, K. B. A., Tan, A. L. T., Yada, S., Sagiraju, S., Bode, P. K. & Barker, N. 2020. Lgr5 Marks Adult Progenitor Cells Contributing to Skeletal Muscle Regeneration and Sarcoma Formation. Cell Rep, 33, 108535.

Lewis, A. E., Vasudevan, H. N., O’neill, A. K., Soriano, P. & Bush, J. O. 2013. The widely used Wnt1-Cre transgene causes developmental phenotypes by ectopic activation of Wnt signaling. Developmental Biology, 379, 229–234.

Liao, J., Huang, Y., Wang, Q., Chen, S., Zhang, C., Wang, D., Lv, Z., Zhang, X., Wu, M. & Chen, G. 2022. Gene regulatory network from cranial neural crest cells to osteoblast differentiation and calvarial bone development. Cell Mol Life Sci, 79, 158.

Madisen, L., Zwingman, T. A., Sunkin, S. M., Oh, S. W., Zariwala, H. A., Gu, H., Ng, L. L., Palmiter, R. D., Hawrylycz, M. J., Jones, A. R., Lein, E. S. & Zeng, H. 2010. A robust and high-throughput Cre reporting and characterization system for the whole mouse brain. Nat Neurosci, 13, 133–40.

Manzanares, M. C., Goret-Nicaise, M. & Dhem, A. 1988. Metopic sutural closure in the human skull. Journal of anatomy, 161, 203–215.

Maruyama, T., Mirando, A. J., Deng, C. X. & Hsu, W. 2010. The balance of Wnt and Fgf signaling influences mesenchymal stem cell fate during skeletal development. Sci Signal, 3, ra40.

Merrill, A. E., Sarukhanov, A., Krejci, P., Idoni, B., Camacho, N., Estrada, K. D., Lyons, K. M., Deixler, H., Robinson, H., Chitayat, D., Curry, C. J., Lachman, R. S., Wilcox, W. R. & Krakow, D. 2012. Bent bone dysplasia-FGFR2 type, a distinct skeletal disorder, has deficient canonical Fgf signaling. Am J Hum Genet, 90, 550–7.

Nagayama, T., Okuhara, S., Ota, M. S., Tachikawa, N., Kasugai, S. & Iseki, S. 2013. FGF18 accelerates osteoblast differentiation by upregulating Bmp2 expression. Congenit Anom (Kyoto), 53, 83–8.

Ng, A., Tan, S., Singh, G., Rizk, P., Swathi, Y., Tan, T. Z., Huang, R. Y., Leushacke, M. & Barker, N. 2014. Lgr5 marks stem/progenitor cells in ovary and tubal epithelia. Nat Cell Biol, 16, 745–57.

Ohbayashi, N., Shibayama, M., Kurotaki, Y., Imanishi, M., Fujimori, T., Itoh, N. & Takada, S. 2002. FGF18 is required for normal cell proliferation and differentiation during osteogenesis and chondrogenesis. Genes & development, 16, 870–879.

Pfaff, M. J., Xue, K., Li, L., Horowitz, M. C., Steinbacher, D. M. & Eswarakumar, J. V. P. 2016. FGFR2c-mediated Erk–Mapk activity regulates coronal suture development. Developmental Biology, 415, 242–250.

Pryce, B. A., Brent, A. E., Murchison, N. D., Tabin, C. J. & Schweitzer, R. 2007. Generation of transgenic tendon reporters, Scxgfp and Scxap, using regulatory elements of the scleraxis gene. Developmental Dynamics, 236, 1677–1682.

Quarto, N., Behr, B. & Longaker, M. T. 2010. Opposite spectrum of activity of canonical Wnt signaling in the osteogenic context of undifferentiated and differentiated mesenchymal cells: implications for tissue engineering. Tissue engineering. Part A, 16, 3185–3197.

Reardon, W., Winter, R. M., Rutland, P., Pulleyn, L. J., Jones, B. M. & Malcolm, S. 1994. Mutations in the fibroblast growth factor receptor 2 gene cause Crouzon syndrome. Nature Genetics, 8, 98–103.

Roybal, P. G., Wu, N. L., Sun, J., Ting, M.-C., Schafer, C. A. & Maxson, R. E. 2010. Inactivation of Msx1 and Msx2 in neural crest reveals an unexpected role in suppressing heterotopic bone formation in the head. Developmental Biology, 343, 28–39.

Sahar, D. E., Longaker, M. T. & Quarto, N. 2005. Sox9 neural crest determinant gene controls patterning and closure of the posterior frontal cranial suture. Developmental Biology, 280, 344–361.

Shi, F., Kempfle, J. S. & Edge, A. S. 2012. Wnt-responsive Lgr5-expressing stem cells are hair cell progenitors in the cochlea. J Neurosci, 32, 9639–48.

Stuart, T., Butler, A., Hoffman, P., Hafemeister, C., Papalexi, E., Mauck, W. M., III, Hao, Y., Stoeckius, M., Smibert, P. & Satija, R. 2019. Comprehensive Integration of Single-Cell Data. Cell, 177, 1888–1902.e21.

Sugimoto, Y., Takimoto, A., Hiraki, Y. & Shukunami, C. 2013. Generation and characterization of ScxCre transgenic mice. genesis, 51, 275–283.

Ten Berge, D., Brugmann, S. A., Helms, J. A. & Nusse, R. 2008. Wnt and Fgf signals interact to coordinate growth with cell fate specification during limb development. Development, 135, 3247–57.

Wang, Y., Dong, Z., Yang, R., Zong, S., Wei, X., Wang, C., Guo, L., Sun, J., Li, H. & Li, P. 2022. Inactivation of Ihh in Sp7-Expressing Cells Inhibits Osteoblast Proliferation, Differentiation, and Bone Formation, Resulting in a Dwarfism Phenotype with Severe Skeletal Dysplasia in Mice. Calcif Tissue Int, 111, 519–534.

Wang, Y., Xiao, R., Yang, F., Karim, B. O., Iacovelli, A. J., Cai, J., Lerner, C. P., Richtsmeier, J. T., Leszl, J. M., A., H. C., Yu, K., Ornitz, D. M., Elisseeff, J., Huso, D. L. & Jabs, E. W. 2005. Abnormalities in cartilage and bone development in the Apert syndrome FGFR2 +/S252w mouse. Development, 132, 3537–3548.

Wilkie, A. O., Slaney, S. F., Oldridge, M., Poole, M. D., Ashworth, G. J., Hockley, A. D., Hayward, R. D., David, D. J., Pulleyn, L. J., Rutland, P. &, Et Al. 1995. Apert syndrome results from localized mutations of FGFR2 and is allelic with Crouzon syndrome. Nat Genet, 9, 165–72.

Xu, Z., Wang, W., Jiang, K., Yu, Z., Huang, H., Wang, F., Zhou, B. & Chen, T. 2015. Embryonic attenuated Wnt/β-catenin signaling defines niche location and long-term stem cell fate in hair follicle. Elife, 4, e10567.

Yin, L., Du, X., Li, C., Xu, X., Chen, Z., Su, N., Zhao, L., Qi, H., Li, F., Xue, J., Yang, J., Jin, M., Deng, C. & Chen, L. 2008. A Pro253Arg mutation in fibroblast growth factor receptor 2 (Fgfr2) causes skeleton malformation mimicking human Apert syndrome by affecting both chondrogenesis and osteogenesis. Bone, 42, 631–643.

Yoshida, T., Vivatbutsiri, P., Morriss-Kay, G., Saga, Y. & Iseki, S. 2008. Cell lineage in mammalian craniofacial mesenchyme. Mech Dev, 125, 797–808.

Yu, K., Xu, J., Liu, Z., Sosic, D., Shao, J., Olson, E. N., Towler, D. A. & Ornitz, D. M. 2003. Conditional inactivation of Fgf receptor 2 reveals an essential role for Fgf signaling in the regulation of osteoblast function and bone growth. Development, 130, 3063–74.

