## Supplemental Table 1 for "FGF Signaling Regulates Development of the Anterior Fontanelle"

| Target | Catalog # | Application | Concentration |
| --- | --- | --- | --- |
| *Fgf18* | ACD 495421 | RNAscope *in situ* | 1:50 |
| *Fgfr2* | ACD 443501-C2 | RNAscope *in situ* | 1:50 |
| *Lgr5* | ACD 312171-C2 | RNAscope *in situ* | 1:50 |
| *Lgr6* | ACD 404961 | RNAscope *in situ* | 1:50 |
| *Scx* | ACD 439981 | RNAscope *in situ* | 1:50 |
| *Sox9* | ACD 401051 | RNAscope *in situ* | 1:50 |
| *Sp7* | ACD 403401-C3 | RNAscope *in situ* | 1:50 |
| *Tnmd* | ACD 430531 | RNAscope *in situ* | 1:50 |
| *Wif1* | ACD 412361-C3 | RNAscope *in situ* | 1:50 |
| Opal 570 | Akoya Biosciences FP1488001KT | RNAscope *in situ* | 1:1000 |
| Opal 620 | Akoya Biosciences FP1495001KT | RNAscope *in situ* | 1:1000 |
| Opal 690 | Akoya Biosciences  FP1497001KT | RNAscope *in situ* | 1:1000 |
| Cleaved Caspase 3 (rabbit) | CST 9661 | Immunofluorescence | 1:400 |
| Runx2 (rabbit) | CST 12556 | Immunofluorescence | 1:400 |
| Sox9 (rabbit) | Novus NBP1-85551 | Immunofluorescence | 1:400 |
| Anti-Rabbit IgG Alexa Fluor 568 (goat) | ThermoFisher Scientific A-11036 | Immunofluorescence | 1:200 |

**Supplemental Table 1:** List of in situ probes and antibodies used in the study.
