## Supplementary figures and images for "FGF Signaling Regulates Development of the Anterior Fontanelle"

### Supplemental Figure 1

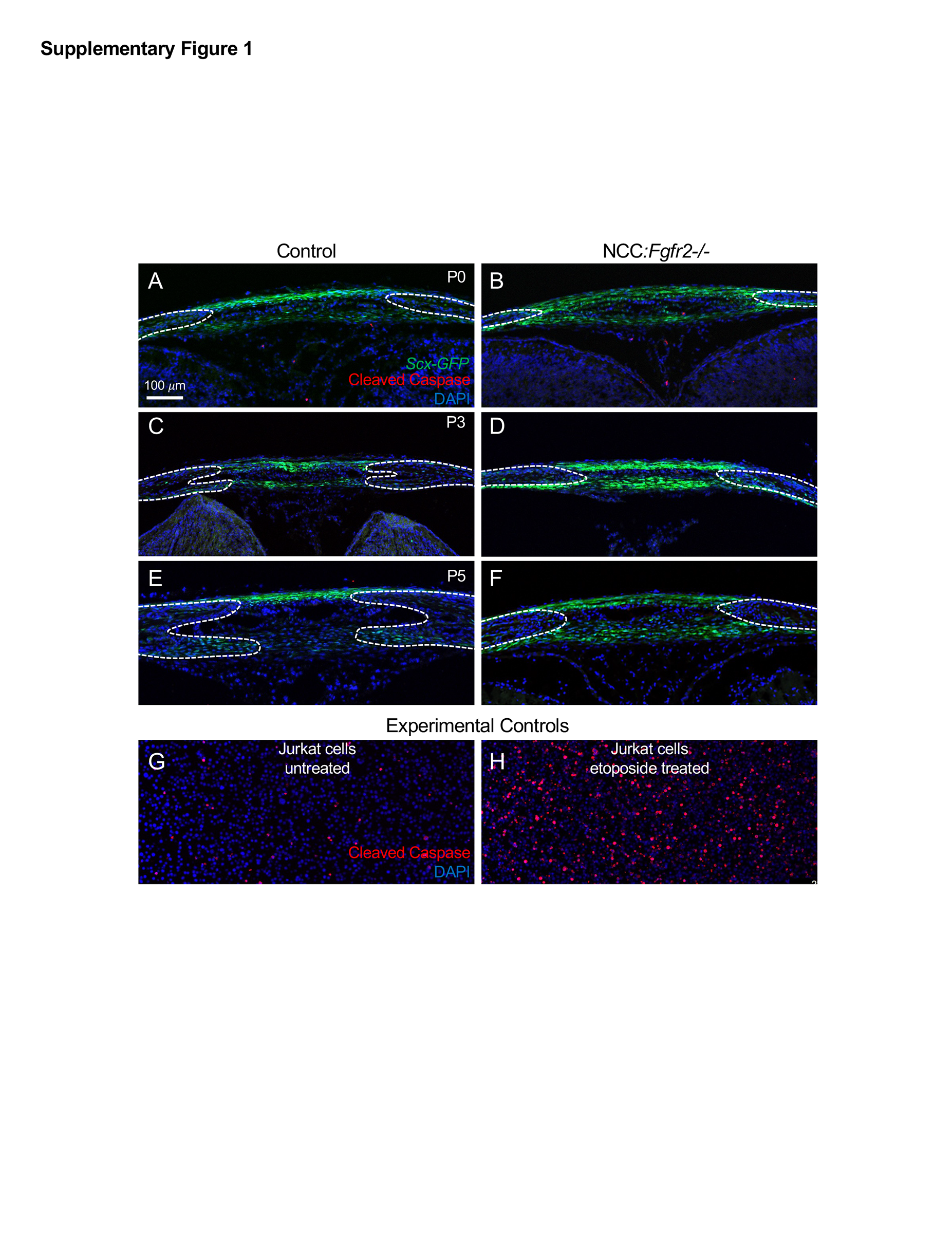
