## Supplementary material for "FGF Signaling Regulates Development of the Anterior Fontanelle": SF1 Legend

**Supplemental Figure Legend**

**Supplemental Figure 1: Cleaved caspase staining shows no significant levels of apoptosis in the developing AF of control or NCC-*Fgfr2^-/-^* mice.** Immunofluorescence staining of cleaved caspase at P0 (A,B), P3 (C,D), and P5 (E,F) shows no significant levels of apoptosis in either NCC-*Fgfr2^-/-^* or littermate control mice (n=3 pairs for each stage). Antibody viability was confirmed with etoposide treated and untreated control slides (CST 8104S) (G,H).
